# AI-assisted isolation of bioactive Dipyrimicins from *Amycolatopsis azurea* and identification of its corresponding *dip* biosynthetic gene cluster

**DOI:** 10.1101/2025.03.21.644653

**Authors:** Christine Mae F. Ancajas, Isra E. Shuster, Allison S. Walker

## Abstract

One of the major challenges in natural product discovery is the prioritization of compounds with useful activities from microbial sources. In particular, this is a challenge in genome mining for novel natural products, where the structures and activities of compounds produced by bioinformatically identified and uncharacterized biosynthetic gene clusters remain unknown. Here, we utilize a machine learning model to predict the antibacterial activity of a natural product from its biosynthetic gene cluster (BGC). We prioritized the strain *Amycolatopsis azurea* DSM 43854 which was predicted by machine learning to have the capacity to produce multiple natural products with antibacterial activity. Together with bioactivity-guided fractionation, we isolated dipyrimicins A and B from *Amycolatopsis azurea* DSM 43854 and, for the first time, linked them to their BGC. This *dip* BGC was predicted by our model to encode a product with 75% antibacterial probability and shares only 40-52% similarity with previously characterized BGCs. We confirmed the antimicrobial properties of the dipyrimicins against a few test strains and identified key tailoring enzymes, including an O-methyltransferase and amidotransferase, that differentiated them from other related 2,2’-bipyridine biosynthetic pathways. Importantly, As the *dip* BGC was not in the training set of the model, our results demonstrate the ability of the model to generalize beyond its training set and the potential of machine learning to accelerate novel bioactive natural product discovery and deorphanization of biosynthetic gene clusters.

## Introduction

With the rise of antibiotic resistance, it has become increasingly important to find novel antibiotic compounds. Natural products, also known as secondary metabolites, are small molecules produced by bacteria, fungi, and plants and serve as a promising source of bioactive molecules. Many natural products have antimicrobial, antifungal, and cytotoxic properties making them useful for the treatment of infectious disease and cancer. Currently, the majority of antibiotic compounds used therapeutically are natural products or natural product derivatives. ^1^ As such, it is important to continue exploring the bioactivity of natural products because this can lead to the discovery of novel antibiotic compounds to combat the spread of antimicrobial resistant pathogens.^2^

In bacteria, the genetic information for natural product biosynthesis is grouped together in what is known as biosynthetic gene cluster (BGC), with each BGC encoding for the enzymes responsible for natural product synthesis. While previous drug discovery efforts have showcased the bioactive potential of natural products, the classical methods used face recurring challenges such as the rediscovery of known compounds and difficulty prioritizing which BGCs are most promising.^3^ We propose that machine learning (ML) techniques can predict which BGCs produce bioactive natural products, leading to more efficient bioactive natural product discovery. Here, we use a method we previously developed to predict the bioactivity of natural products produced by BCGs.^4^ Our previous work has shown that rare actinomycetes and underexplored genera are promising candidates for antibiotic discovery.^5^

To test the accuracy of the ML model in tandem with experimental methods, we worked with *Amycolatopsis azurea* DSM 43854 as a promising candidate for antibiotic production. *A. azurea* DSM 43854 belongs to a rare genera of actinomycete with great potential for novel antibiotic discovery.^6^ Over 25 orphan BGCs with less than 75% similarity to characterized BGCs were predicted to produce an antibiotic compound. In this study, we used these predictions to guide our exploration of natural products produced by *A. azurea* DSM 43854. We isolated and purified antibiotic compounds produced by *A. azurea* DSM 43854, which we identified as dipyrimicin A and B. Dipyrimicin A and B were first identified and isolated from *Amycolatopsis* sp. K16-0194 in 2018 as antibiotic compounds, but the BGC of origin was unknown.^7^ Therefore, the dipyrimicin BGC is not present in the training data used to train the machine learning model used in this study. Our work serves as a proof of concept for machine learning guided discovery of antibiotic natural products, as the algorithm predicted that the BGC responsible for the production of dipyrimicin A and B would produce an antibiotic compound. Further *in silico* analysis identified the biosynthetic origin of dipyrimicin A and B, as well as specific enzymes responsible for the synthesis of specific functional groups within the compound.

## Results and Discussion

### Genome mining and ML model-guided screening reveals promising novel BGCs

Secondary metabolites remain as an important source of bioactive compounds. However, isolating active compounds is often hindered by the uncertainty of whether a given bacterial strain can produce metabolites with useful activities. To assess potential microbial sources for active compounds, specifically with antibiotic activity, we utilized a machine learning workflow, which uses annotations from antiSMASH 5 and RGI 5, to predict the antibiotic activity of BGCs within a given bacterial genome.^4,8,9^ We considered BGC activity prediction of 50% or greater as the “active” threshold as this represents a greater than even likelihood of producing an active compound. Based on this criteria, *A. azurea* DSM 43854 was identified as a promising candidate. First, antiSMASH analysis revealed that the genome of *A. azurea* DSM 43854 encodes 35 BGCs, many of which were <75% similar to known BGCs and classified across diverse natural product classes, including NRPS, PKS, RiPPs, terpenes, and more (**Figure 1, Table S1)**. Moreover, of the 35 BGCs, 25 BGCs meet or exceed the “active” threshold for antibacterial activity (**Figure 1, Table S2**).

**Figure 1.**
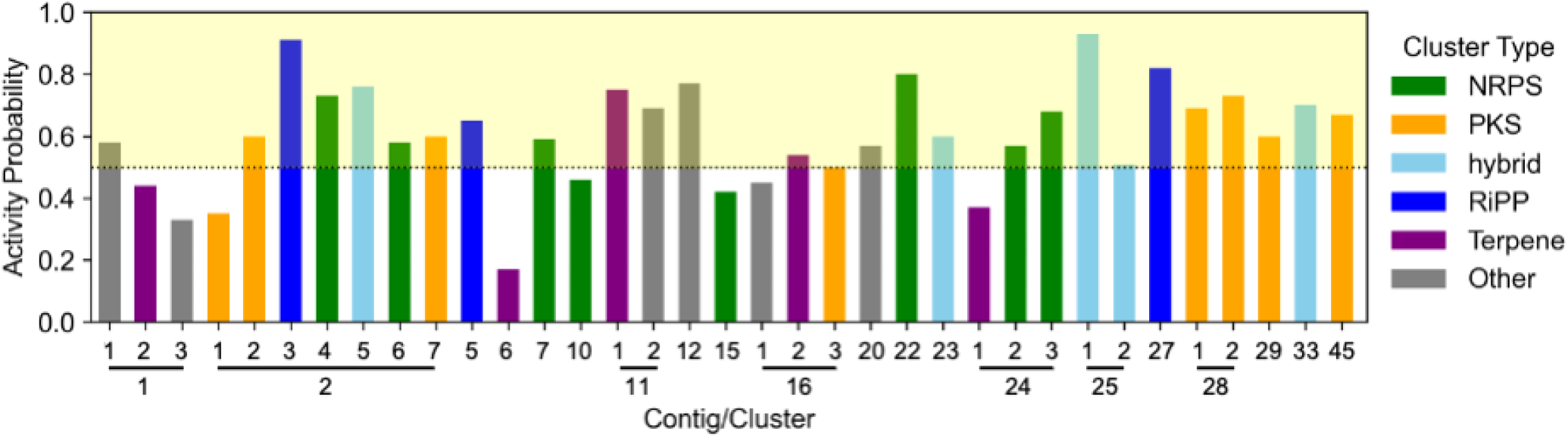
ML-based antibiotic activity prediction for each BGC in *Amycolatopsis azurea* DSM 43854. A probability threshold of >50% indicates higher antibiotic activity potential. Each BGC is color-coded by its antiSMASH predicted natural product class: green for NRPS, orange for PKS, light-blue for hybrid, dark-blue for RiPPs, purple for terpene, and gray for other. Clusters located on the same contig are grouped together, as indicated by the lines.

To the best of our knowledge, only two families of compounds have been isolated from *Amycolatopsis azurea* DSM 43854: 1) azureomycins A and B ^10–12^ 2) octacosamicins A and B.^13,14^ Azureomycins, discovered in 1979, was reported to have peptidoglycan synthesis inhibition activity and physico-chemical properties resembling glycopeptides suggest that they are putative novel glycopeptides, although their exact structures remain uncharacterized.^10–12,15^ AntiSMASH 5 analysis identified contig 25.1 as the only glycopeptide-encoding BGC (92% similar to keratinimicins, **Table S1**) which suggests this cluster as the likely source of the uncharacterized azureomycins, aligning with both activity prediction (93% activity probability, **Figure 1**) and previous reported data. On the other hand, Liao et al. induced the production of antifungal polyketide, octacosamicin A, by sub-inhibitory concentrations of streptomycin and bioinformatically linked it its BGC, a proposed combination of PKS clusters 16.3 and 2.1.^14^ The presence of several other orphan BGCs in *A. azurea* DSM 43854, such as clusters in contigs 2, 11, and 28, each with a high likelihood of producing diverse bioactive compounds (**Figure 1**), highlights this strain as an excellent starting point for our ML-prioritized bioactive natural product discovery efforts.

### Identification of active compounds

After identifying *A. azurea* DSM 43854 as a potential candidate for producing antibiotics, we proceeded to experimentally validate the BGC predictions by isolating bioactive metabolites. Initially, the strain was cultured under various media conditions followed by liquid-liquid and solid-phase extractions. Crude extracts obtained were screened for antibacterial activity against *Bacillus spizizenii* ATCC 6633 by observing zones of inhibition. Extracts of cultures grown in ISP4 agar plates showed a notably higher zone of inhibition. We then proceeded to perform bioactivity-guided fractionation until individual active metabolites were isolated.

Of the fractions with notable antibiotic activity, two components – herein, compound **1**, eluting first, and compound **2** – exhibited prominent UV absorption spectra at around 205, 240, 270, and 320 nm (**Figure S1**), characteristic of aromatic chromophores. High resolution ESI-MS analysis for **1** identified an m/z of 247.0708 [M+H]^+^ and m/z 246.0871 [M+H]^+^ for **2** (**Figure S2**). To note, despite maintaining similar growth conditions, production of compound 2 was less reproducible across independent cultures, suggesting that its production may be influenced by subtle variations in growth conditions. The relative similarity in their retention time and UV absorption spectra suggest these are closely related compounds. Based on previous reports, these represent new metabolites isolated from *A. azurea* DSM 43854 as the observed m/z values differ from those of the known octacosamicins (m/z 625 and 639)^13,14^ and azureomycins (m/z >850);^11^ to note, under our specific fermentation and extraction conditions, no m/z values corresponding to these two metabolites were detected.

To confirm the structures, comprehensive NMR experiments were performed. ^1^H NMR revealed aromatic signals (δ_H_ 7.75 – 8.9), correlating with the UV absorption data, and a methoxy group (around δ_H_ 4.01 and δ_C_ 55.2) in **1** and **2** (**Figure 2, Table 1**). Furthermore, detailed 2D NMR (COSY, HSQC, and HMBC) analysis confirmed the aromatic presence as a 2,2’-bipyridine (2,2’-BP) core (**Figure S3-S10**), which aided in our analysis. Database and literature search identified similarities with the well-characterized 2,2’-BP-containing natural products collismycins and caerulomycins.^16^ Additionally, the reported NMR by Izuta et al.^7^ aligns with our observed connectivities of the hydroxyl group on C3’, methoxy group on C4’, aromatic proton H5’, and carbonyl group on C7’ on the more substituted ring. The 1 m/z difference between compound **1** and **2** can be attributed to presence of the hydroxyl or amine functional group on C7’, respectively (**Figure 2**). This is in agreement with the chemical formulas C_12_H_10_N_2_O_4_ (calcd. 247.0719) and C_12_H_11_N_3_O_3_ (calcd. 246.0879) as well as the observed m/z values reported both in this study (**Figure S2**) and by Izuta et al.^7^ Therefore, in accordance with the previous data, compound **1** and **2** from the active fraction are, hereafter, confirmed as dipyrimicin A and dipyrimicin B, respectively.

**Table 1.**
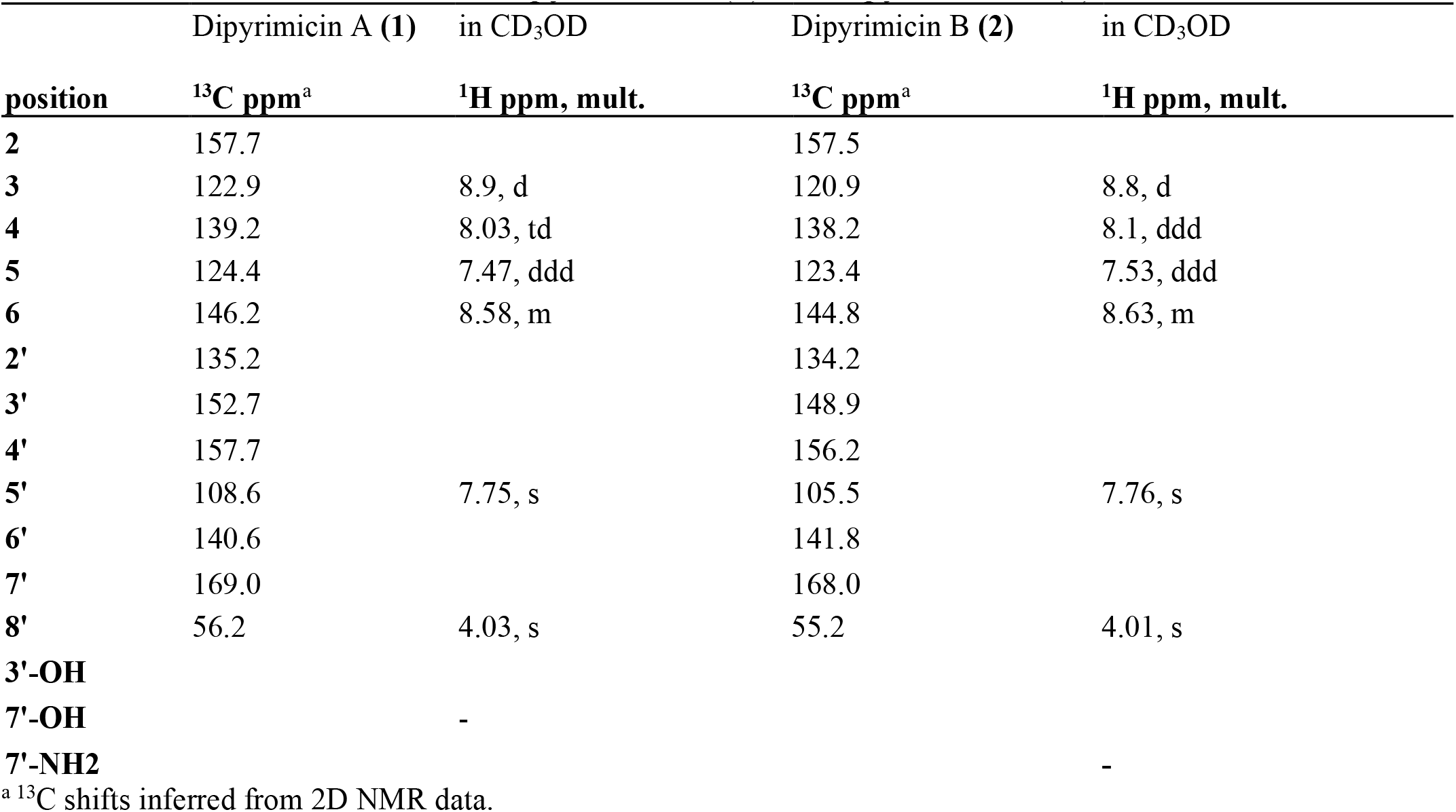
Chemical Shift Data for dipyrimicin A (**1**) and dipyrimicin B (**2**).

**Figure 2.**
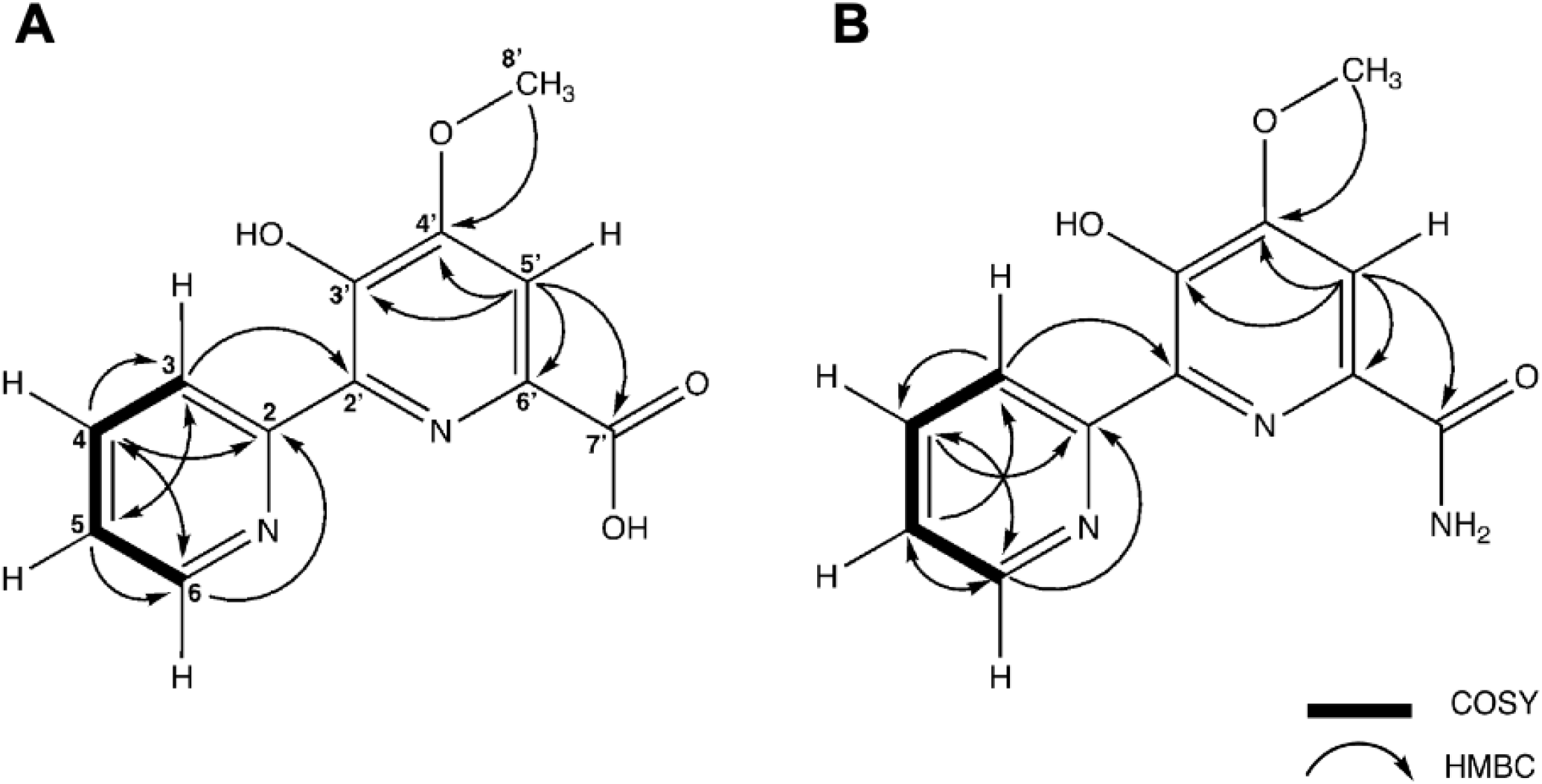
Key COSY and HMBC correlation observed for **(A)** dipyrimicin A (compound **1**) and (**B**) dipyrimicin B (compound **2**) isolated from *A. azurea* DSM 43854.

### Bioinformatic analysis of Dipyrimicin BGC and biosynthesis

Having confirmed the production and initial activity of dipyrimicins A (**1**) and B **(2)** by *A. azurea* DSM 43854, we analyzed the genome to link the isolated compounds to their biosynthetic origin. Dipyrimicins, first reported by Izuta et al. in 2018, were originally isolated from a closely related strain, *Amycolatopsis* sp. K16-0194.^7^ Since then, dipyrimicin has also been reported from *Streptomyces thioluteus*,^17^ along with other 2,2’-BP analogs from various strains.^18^ However, high-quality genome assemblies and the BGCs responsible for these compounds have yet to be reported. Moreover, identification of this uncharacterized BGC is of particular interest as it was absent in the training data used to develop the ML model applied in this study, which allows us to evaluate whether the model can generalize beyond its training set. Here, to identify the dipyrimicin BGC (*dip*), we refined our search to the 25 BGCs with high activity probabilities. Among these, we identified cluster 2.5 (76% antibacterial activity probability) as the putative *dip* BGC **(Figure 1**). This cluster shares 52% and 40% similarity with the BGCs for caerulomycin A (*cae*) and collismycin A (*col*), respectively.

As the dipyrimicins share the 2,2’-BP core of the caerulomycins and collismycins, we examined the putative *dip* BGC for the presence of genes conserved in their respective pathways. Comparison of the annotated genes within the putative *dip* BGC to those of *cae* and *col* BGCs indicated that all of the genes necessary for formation of a 2,2’-BP core were present, ranging from 25% to 79% identity (**Table S3**). Specifically, *dip11-17* shows close homology to the conserved *cae/col A1-A4, P1-P2*, and *B1* genes which form the 2,2’-bipyridine-L-leucine intermediate, as well as the transporter genes *cae/col H1-H2* (**Figure 3)**.^19,20^ Distant homologs of the pathway-specific regulators, *cae/colI1-I2*, were also present.^21^ However, the *dip* BGC lack the genes (*cae/col B2, F, C, and caeB5*) likely involved in forming the oxime group characteristic of the caerulomycins and collismycins (**Table S3**).^22–25^ Subsequently, we identified notable *dip* genes, including those encoding an O-methyltransferase and amidotransferase, which differ from either of the known *cae* and *col* pathways. These enzymes are hypothesized to decorate the 2,2’-BP core to afford the dipyrimicins (**Figure 4**). While the functional groups installed are not entirely unique to the dipyrimicins, identifying the enzymes responsible for these functionalities supports the assignment of the *dip* BGC and provides new insights into the biosynthetic pathways within this family of compounds.

**Figure 3.**
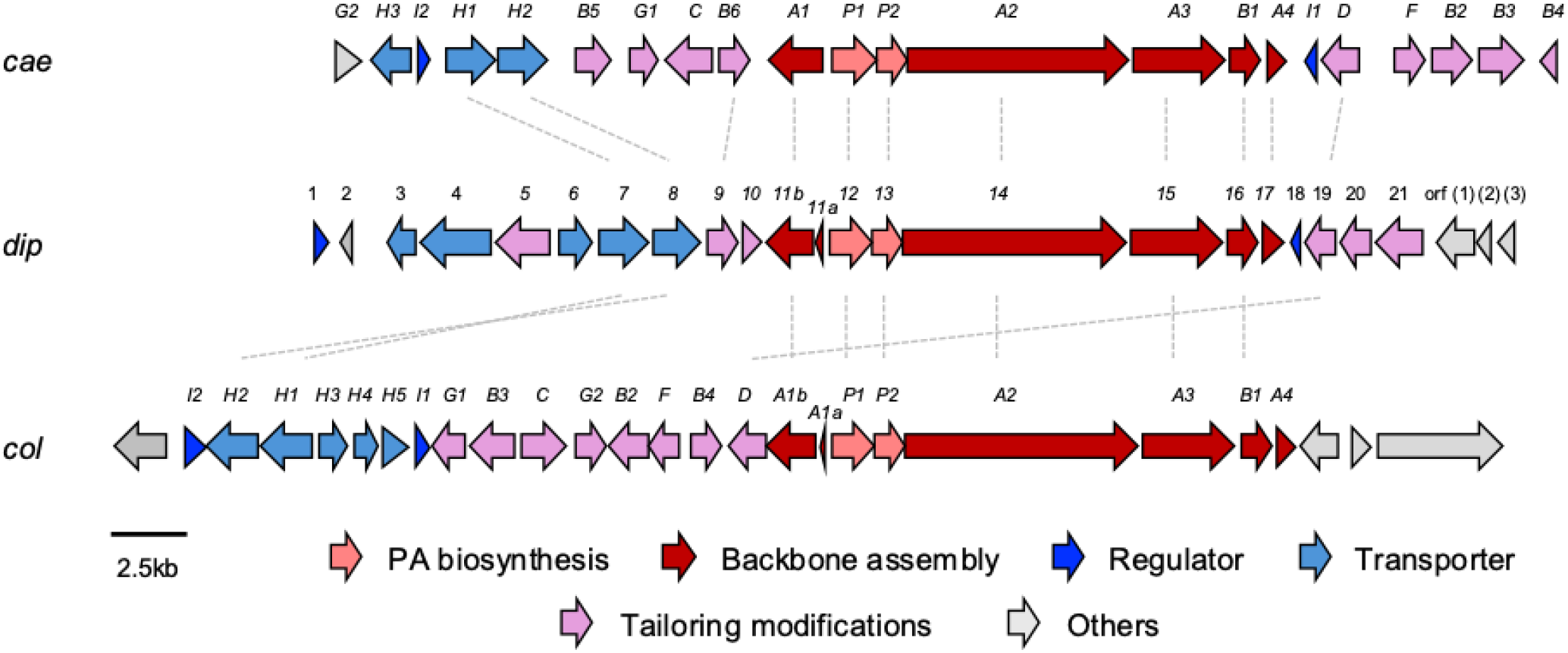
Alignment of the **(A)** caerulomycin A BGC (*cae*) from *Actinoalloteichus* sp. WH1-2216-6, **(B)** putative dipyrimicin A and B BGC (*dip*) from *A. azurea* DSM 43854, and **(C)** collismycin A BGC (*col*) from *Streptomyces sp*. CS40. Alignments with >50% identity are linked by dashed lines. Standard nomenclature for *cae* and *col* by the Liu group were adapted.^26^

**Figure 4.**
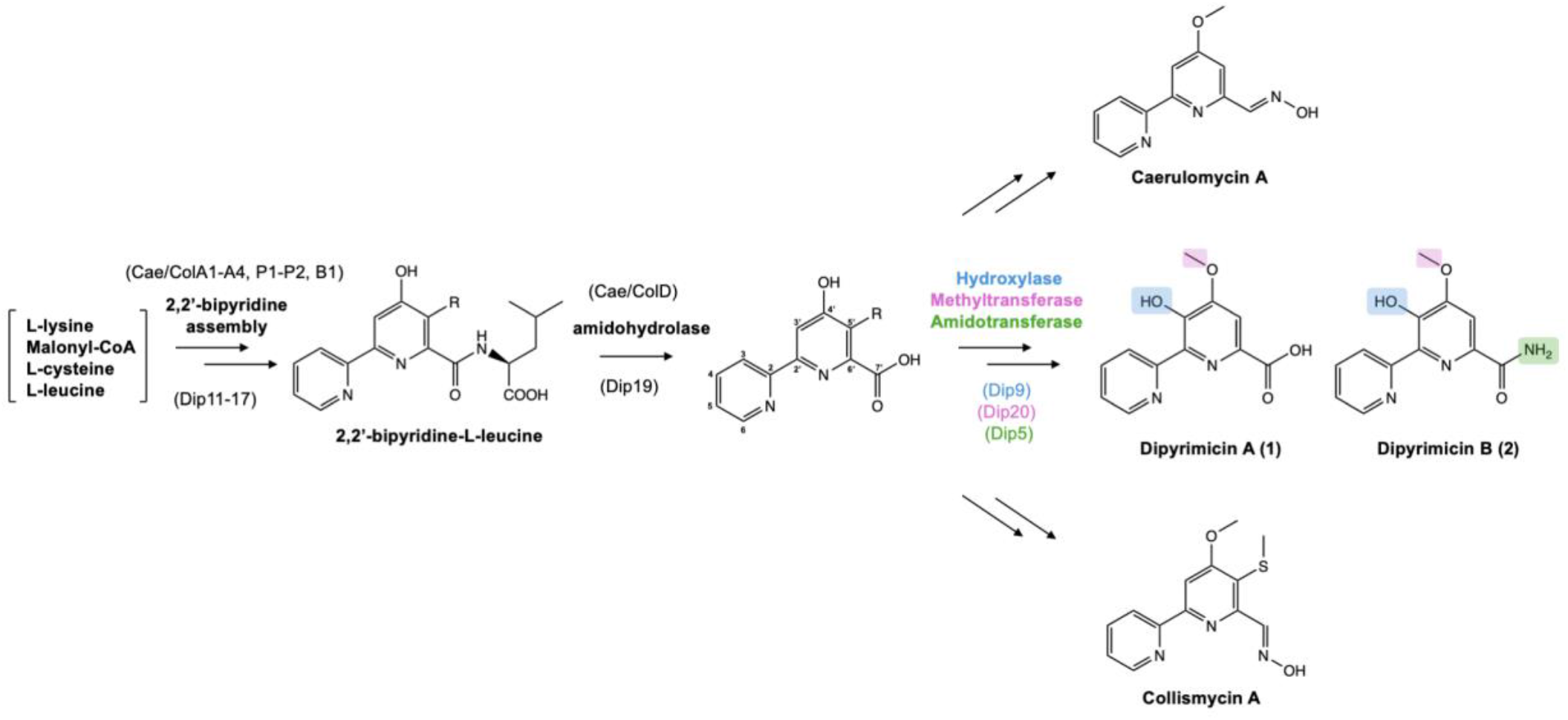
The biogenesis of dipyrimicins follows a similar pathway to caerulomycins and collismycins through the formation of a 2,2’-bipyridine (2,2’-BP) scaffold. Variation in tailoring modifications differentiate them, such as the oxime group in caerulomycin A and collismycin A. Dipyrimicins A (**1**) and B (**2**) are hydroxylated at C3’ and O-methylated at C4’ while dipyrimicin B is further amidated at C7’.

First, the presence of *dip19*, which encodes an amidohydrolase with 73% and 61% identities to CaeD (83% similarity) and ColD (73% similarity), respectively, suggests that Dip19 catalyzes the same hydrolysis reaction, converting the 2,2’-BP-L-leucine intermediate into the carboxylated 2,2’-BP retained in dipyrimicin A **(Figure 4)**.^27^ The other tailoring gene with relative homology within the *cae* BGC is *dip9* which is predicted to encode a NAD(P)/FAD-dependent oxidoreductase (65% identity, 71% similarity) and shows the highest sequence homology to *caeB6* (54% identity, 66% similarity). To note, no homology of this gene is found in the *col* BGC and, as expected, there are no reported C3’-hydroxylated collismycins. In caerulomycins, CaeB6 has been experimentally implicated to catalyze the oxygenation at C3’.^28^ Therefore, it is likely that Dip9 putatively hydroxylates C3’ in the dipyrimicins as well.

Similar to several caerulomycins and collismycins, dipyrimicins are O-methylated at C4’. In the *cae* and *col* pathways, this formation is processed by O-methyltransferases CaeG1 and ColG1, respectively.^22,29^ Previous experimental studies by Chen et al. on the *cae* pathway revealed a competition between the CaeB6 C3’-hydroxylation and CaeG1 C4’-OH methylation of caerulomycin H, an oxime-containing C4’-O-demethylated 2,2’-BP intermediate. The main route leading to caerulomycin A involves direct C4’-OH methylation where C3’ remains unmodified. In the minor route, CaeB6 hydroxylates C3’ before CaeG1 C4’-OH methylation.^28^ Within the putative *dip* BGC, a hypothetical protein, Dip20, shares the highest similarity to SAM-dependent methyltransferases (65% identity and 74% similarity) but exhibits low homology to CaeG1 (25% identity, 42% similarity) and ColG1 (26% identity, 44% similarity). BlastP analysis from MIBiG reveals that Dip20 has close homology to QbsL (43% identity, 63% similarity) in the siderophore quinolobactin pathway.^30^ QbsL is a hybrid protein with an AMP-dependent ligase and synthetase domain (N-terminal) and a methylase domain (C-terminal), facilitating the carboxylic acid activation and O-methylation of a hydroxyl group, respectively, on a precursor xanthurenic acid.^31^ We propose that the dipyrimicin pathway adopts the minor route in caerulomycin biosynthesis as its primary pathway where Dip9 hydroxylates C3’, followed by C4’-OH methylation by Dip20. The resulting formation of the C4’-methoxy group likely serves as a protecting group, preventing additional modifications on the hydroxyl group that could potentially lead to 2,2’-BP glycosylated cyanogrisides and other derivatives.

The amido group in dipyrimicin B (**2**) has been observed in a few 2,2’-BP metabolites, as synthetic derivatives (eg. caerulomycinamide)^17,32,33^, shunt products (eg. collismycin DS)^23^, and/or isolated from bacterial cultures (eg. caerulomycinamide and saccharobipyrimicin 1)^17,18^ yet no specific enzyme has been linked to its formation. To investigate a possible enzyme for this modification, we examined *dip5*, which encodes a putative amidotransferase with high similarity to glutamine-dependent asparagine synthetases (62% identity, 73% similarity) known to catalyze the ATP-dependent transfer of a glutamine-derived amino group to aspartate.^34^ Additionally, Dip5 shows homology to DacD (51% similarity, 63% identity) and other N-terminal-nucleophilic (Ntn) enzymes involved in conversion of a malonate-equivalent starter unit to a corresponding malonamate in tetracycline family biosynthesis.^35,36^ Based on these similarities, Dip5 may also catalyze an amine transfer at C7’, converting dipyrimicin A into the amidated dipyrimicin B.

While the putative tailoring genes encoding enzymes that modify the 2,2’-BP core to produce the dipyrimicins have been annotated through sequence analysis, the availability of advanced structure prediction technologies, such as AlphaFold2, enables the integration of sequence and structural data to further refine the functional assignments of these tailoring enzymes. Using the open-source implementation of AlphaFold2 in ColabFold,^37^ protein models for these enzymes were generated, and FoldSeek^38^ was used to search the structural space within the Protein Data Bank (PDB) and AlphaFold Database (AFDB). For example, since Dip9 shows high sequence homology to Cae/ColB6, it is no surprise that proteins with similar structures identified by FoldSeek belong in the oxidoreductase superfamily, although the highest sequence identities are only around 36-39% (**Table S4**). Also, the AlphaFold-predicted structure of Dip9 showed strong alignment to the predicted model of CaeB6 and other related hydroxylases (**Figure S11**). Similarly, structural analyses of the Dip20 O-methyltransferase, along with the predicted model of QbsL, confirmed its classification within the SAM methyltransferase superfamily **(Table S5)**, as indicated by the conserved “DxGxGxG” motif for SAM binding (**Figure S12**).^39^ For the putative Dip5 amidotransferase, the closest structural homologs in the AFDB were glutamine-dependent asparagine synthetases, with sequence identity as high as ∼45% (**Table S6**). Structural alignment with related proteins, including the predicted structure of DacD, confirms that Dip5 contains the conserved ATP-binding motif (“SGGLDS”)^40^ and the distinct N-terminal glutaminase and C-terminal synthase domains **(Figure S13)**. Furthermore, similar to the hydroxyamidotransferase TsnB9 (**PDB 7YLZ**), Dip5 possesses inserted regions that differentiate it from the canonical asparagine synthetase, AsnB from *Escherichia coli* (**PDB 1CT9**), potentially allowing it to accommodate the dipyrimicin acid substrate (**Figure S13**).^41^

Lastly, the potential biosynthetic roles of two genes, *dip10* and *dip21*, remain unaccounted for. Dip21 is located adjacent to the putative Dip20 O-methyltransferase and is predicted to function as an AMP-dependent synthetase and ligase (**Table S3**), sharing 48% identity and 64% similarity with QbsL. Notably, the xanthurenic acid processed by QbsL methylates a hydroxyl group meta to a carboxylic acid activated by the ligase, a similar structural arrangement observed in the dipyrimicin acid.^31^ Structural comparisons indicate that the separate AlphaFold models of Dip20 and Dip21 align well with the C-terminal methylase and N-terminal synthetase domains of AlphaFold QbsL structure, respectively, suggesting similar functions (**Figure S11**). FoldSeek search revealed homologs with ligase activity including aminoacyl-AMP synthetase and CoA ligases (**Table S7**). However, their precise roles in dipyrimicin biosynthesis remain unclear. If Dip21 is functional, its role as an activator may be relevant as the intermediate it could form, such as dipyrimicin-AMP, would be highly susceptible to nucleophilic attack. In the quinolobactin biosynthesis, QbsL-mediated carboxyl activation facilitates the subsequent sulfurylation and reduction by QbsCDEK, leading to the formation of a thioacid before a proposed spontaneous hydrolysis regenerates the carboxyl group.^42^ However, homologs of these sulfurylation and reduction enzymes are absent in the *dip* BGC. Furthermore, while the acyl-adenylate intermediate formed could theoretically be targeted by an N-source for amidation, Dip5 has been shown to contain the corresponding domains to catalyze its own adenylation (**Figure S10**) which is typical for most asparagine synthetases,^34^ suggesting a different purpose for Dip21 activation. Meanwhile, Dip10 exhibits the highest sequence similarity to an α/β-hydrolase (67% identity, 76% similarity). Homology searches against MIBiG reveal similarities to the Ntn hydrolase PvdQ from the pyoverdine siderophore biosynthetic pathway (45% similarity, 61% identity), which catalyzes the hydrolysis of a pyoverdine precursor and a fatty acid chain.^43^ Structural homology hits from FoldSeek identified related hydrolases (**Table S8**). If Dip10 functions analogously to these related proteins, it may hydrolyze at the electrophilic center of an intermediate to regenerate the carboxyl group in dipyrimicin A.

Based on sequence and structural analyses, we propose the biosynthetic pathway of dipyrimicins in *Amycolatopsis azurea* DSM 43854 (**Figure 5**). Our hypothesized pathway follows the conserved 2,2’-BP formation but diverges from the known caerulomycin and collismycin pathways after Dip9-mediated C3’-hydroxylation. Instead, dipyrimicin biosynthesis appears to undergo a distinct O-methylation mechanism in which Dip20 O-methyltransferase is adjacent to an Dip21 AMP-ligase. However, whether Dip20 and Dip21 function cooperatively, as in the bifunctional QbsL, remains uncertain. The low sequence and structural homology of Dip20 to Cae/ColG1 may reflect not only differences in substrate specificity or enzymatic promiscuity but could also indicate that Dip20 was acquired from a distinct yet related siderophore biosynthetic pathway. Unlike Dip5, Dip9, and Dip20, where the corresponding enzymatic reactions and modifications in the dipyrimicin biosynthesis are well-supported, the roles of Dip21 and Dip10 remain uncertain as the pathway could be complete without them. To complete the *dip* BGC annotation, *dip3, dip4*, and *dip6* encode ABC and MFS transporters likely involved in the export of the natural product while *dip2* encodes a hypothetical protein with unknown function (**Table S3**). To validate our bioinformatic analyses, experimental studies will be essential to confirm the functional roles especially for the post-tailoring modifying enzymes encoded within the putative *dip* BGC.

**Figure 5.**
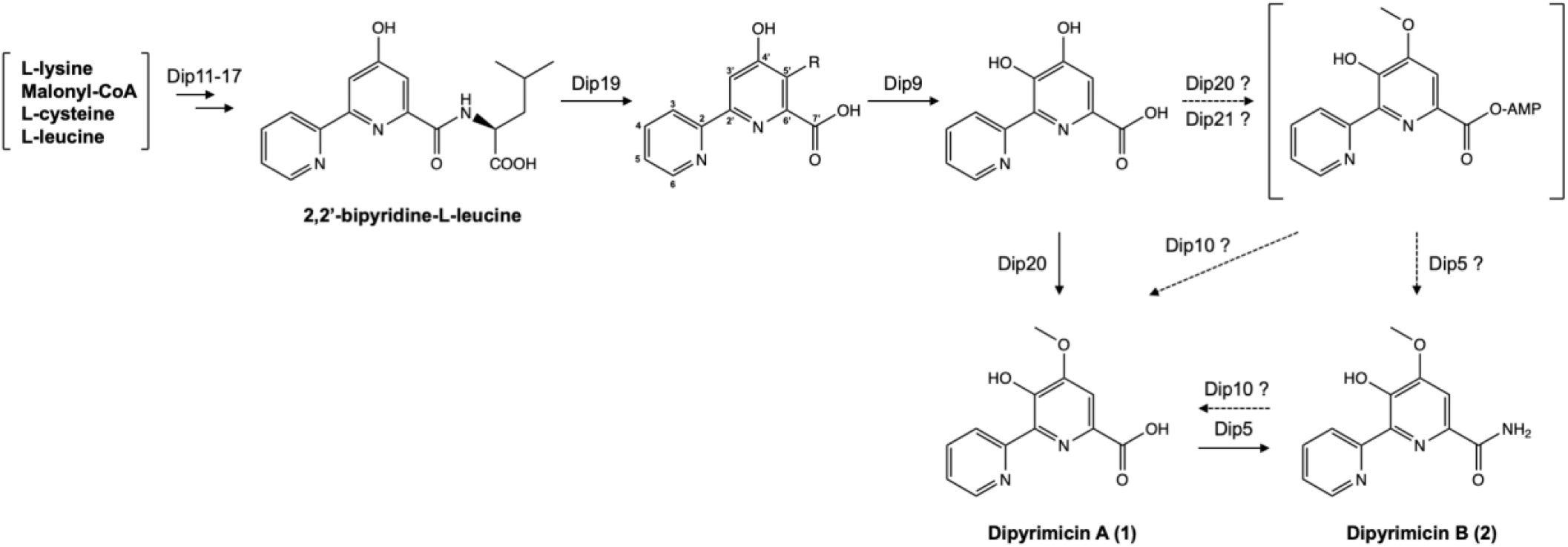
The proposed biosynthesis of dipyrimicin A (**1**) and dipyrimicin B (**2**) in *A. azurea* DSM 43854 follows the caerulomycin and collismycin pathways in which hybrid PKS/NRPS enzymes (Dip11-17) form the 2,2’-bipyridine-*L*-leucine intermediate followed by cleavage of leucine by Dip19 amidohydrolase. A homolog of Cae/ColB5, Dip9, hydroxylates at C3’ and putative Dip20 O-methyltransferase forms the methoxy group at C7’ to form dipyrimicin A which could be further processed by Dip5 amidotransferase to form dipyrimicin B (**2**). An alternative pathway in which Dip21-Dip20, similar to hybrid AMP-ligase and methylase QbsL, may C4’-methylate and activate the carboxylic acid to form the acyl-adenylate intermediate. Dip5 and putative Dip10 hydrolase may function to form the dipyrimicin B and regenerate the carboxylic acid in dipyrimicin A.

### Biological activity

Through bioactivity-guided fractionation and the identification of the *dip* BGC, we confirmed that the ML-predicted BGC produces bioactive metabolites. However, while the ML model predicts bioactivity, it does not quantify potency. Therefore, we further characterize the antimicrobial activity of the dipyrimicins by determining their minimum inhibitory concentrations (MICs) of the dipyrimicins and establishing preliminary structure-activity relationship (SAR) trends.

Previous susceptibility studies using the disk diffusion method reported moderate antimicrobial activity of dipyrimicin A with growth inhibitions of 16-23 mm at doses of 30 µg and 100 µg dose against *Escherichia coli* NIHJ and *Bacillus spizizenii* ATCC 6633. In contrast, dipyrimicin B only exhibited activity at 100 µg against *E. coli* NIHJ.^7^ Here, we confirm the minimum inhibitory concentrations (MIC) of the dipyrimicins via broth microdilution. Dipyrimicin A displayed moderate activity against Gram-positive *B. spizizenii* ATCC 6633 (32 µg/mL) and *Staphylococcus aureus* ATCC 25923 (64 µg/mL), while its activity was reduced against Gram-negative *E. coli* MG1655 ATCC 700926 (128 µg/mL) and *Pseudomonas aeruginosa* (>128 µg/mL). Consistent with previous bioassays, dipyrimicin B displayed weaker activity with partial inhibition at 128 µg/mL for both *B. spizizenii* ATCC 6633 and *E. coli* MG1655 ATCC 700926 (**Table S9**). These findings suggest that Dip5 amidation on C7’ in dipyrimicin B diminishes antibacterial activity. Moreover, these MIC values are relatively weak in comparison to those reported for other 2,2’-BP natural products, which have demonstrated activity as low as 2.5 to 5 µg/mL against *E. coli* and *P. aeruginosa*, respectively, by caerulomycin A,^33^ and 8 ug/mL against methicillin-resistant *S. aureus* (MRSA) by collismycin A.^44^ This discrepancy in antibacterial activity highlights the well-established importance of the oxime group in enhancing antibacterial activity of the 2,2’-BP compounds.

However, presence of the carboxylic and amido group may serve other biological roles beyond antibacterial activity. For example, previous studies on caerulomycin A analogs revealed that conversion of the aldoxime to a carboxylic or amido functional groups preserved their phytotoxic properties. This was attributed to their ability to form complexes with iron.^45^ Indeed, 2,2’-BP compounds are well-documented to have iron-chelating properties and have been implicated in their diverse bioactivities.^16^ For example, collismycin C, originally reported to exhibit poor antibacterial activity,^46^ was later identified as a potent inhibitor of MRSA biofilm formation, which is a property speculated to arise from its iron-chelating ability and the positioning of the hydroxyl group on the 2,2’-BP core.^47^ More recently, Henriquez et al. proposed that these metal-chelating compounds of the 2,2’-BP family may disrupt intracellular ion homeostasis, leading to microbial metabolic stress and eventual growth inhibition.^48^ Thus, further studies could explore additional bioassays to better understand the mechanism of action of dipyrimicins.

### Expanding targeted genome mining with ML

With the successful identification of a bioactive natural product and its previously unreported BGC, we next explored how this approach could be extended to systematically mine other 2,2’-BP-producing strains. By integrating ML-based activity predictions with targeted genome mining of conserved 2,2’-BP core biosynthetic genes, we identified several promising BGCs with >50% activity probability (**Table S10**) in previously unreported strains such as *Nocardia panacis, Micromonospora craniellae*, and *Micromonospora tulbaghiae*, as well as other species within the *Amycolatopsis* and *Streptomyces* genera. Interestingly, we also identified *Streptomyces caatingaensis*, containing the same gene arrangement as the putative *dip* BGC in *A. azurea* DSM 43854, with >50% sequence identity (**Figure 6**). This suggests that the cis-located tailoring Dip20 methyltransferase and Dip5 amidotransferase may not be random insertions but functionally significant tailoring enzymes, as their presence is conserved across different species between *A*. azurea and *S. caatingaensis*.^49^

**Figure 6.**
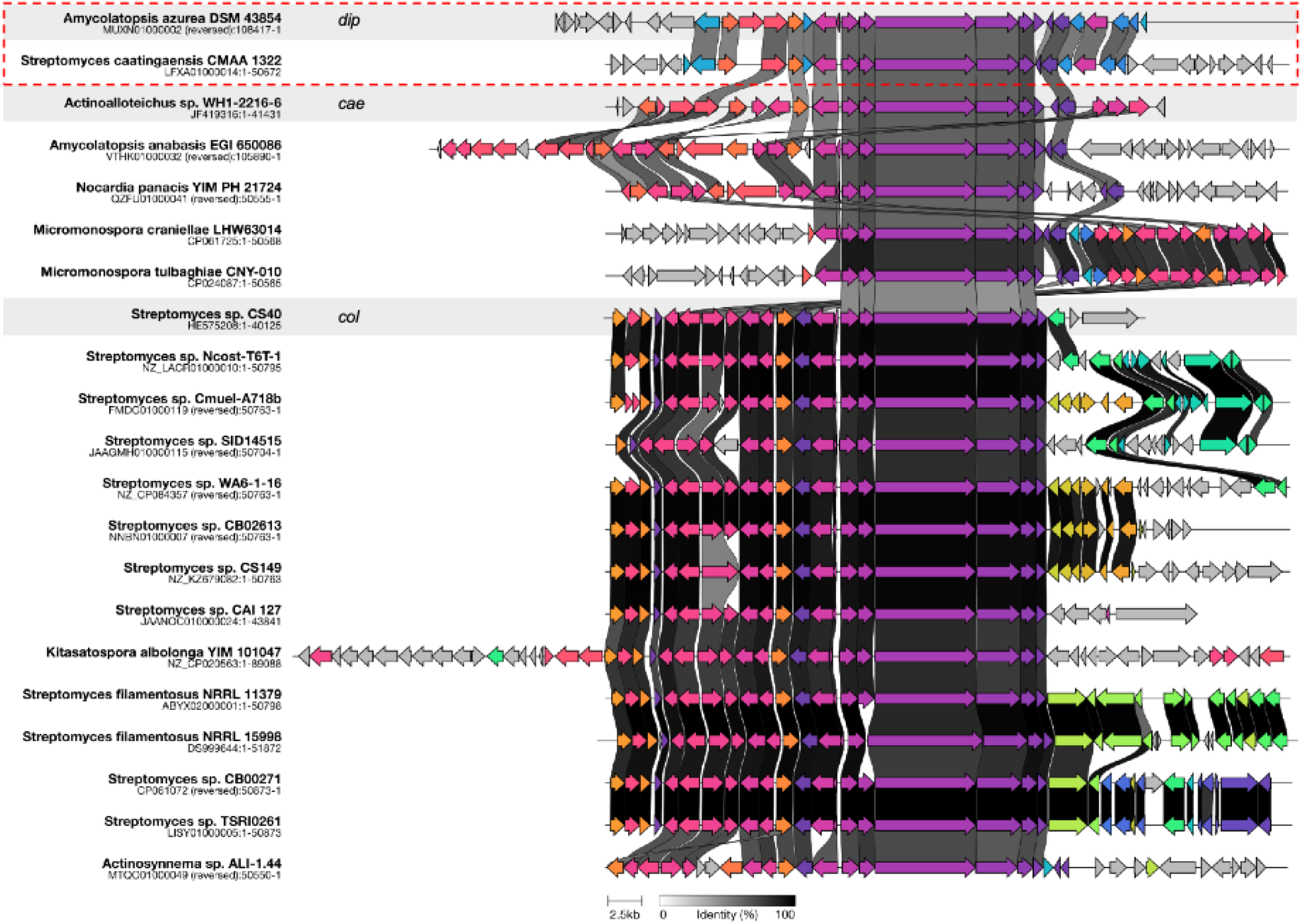
Clinker alignment, accessed through the CAGECAT webserver, of BGCs that are predicted to encode novel 2,2’-BP natural products from unreported bacterial strains.^50^ The clusters highlighted in the gray boxes represent the reference *dip, cae*, and *col* BGCs. The genes within the red dashed box correspond to the putative *dip* BGC found in *Amycolatopsis azurea* DSM 43854 (this paper) and a potential new producer, *Streptomyces caatingaensis* CMAA 1322. Genes that are homologous are color-coded similarly, and greyscale connections between genes further indicate >50% sequence identity.

Our genome mining analysis also highlights the variation in tailoring modifications around the 2,2’-BP core, suggesting further potential for more structural diversity among 2,2’-BP analogs (**Figure 6)**. As demonstrated by the dipyrimicins and other members of the 2,2’-BP family, these tailoring modifications play a crucial role in modulating biological function. This underscores a key limitation in traditional genome mining, which relies primarily on identifying conserved biosynthetic genes ie. the presence of the 2,2’-BP core may not solely be predictive of antimicrobial potential. Therefore, genome-mining approaches could benefit from using our ML method in tandem by considering the overall genes within the BGC, allowing for a more informed prioritization for natural product discovery.

Apart from 2,2’-BP biosynthesis, our ML approach can be integrated with targeted genome mining to prioritize other natural product scaffolds and bioactive features,^51^ including activity-associated genes.^4^ However, a key challenge in enhancing ML-based predictions is the limited availability of high-quality data. We expect that as we are able to grow our training set, the accuracy of our machine learning model will improve, further accelerating the discovery of novel bioactive natural products. Therefore, it is beneficial for the broader natural product community for genome assemblies, BGC annotations, and the activities, including lack of activity, of the linked natural products to be reported in publications and shared public databases such as MIBiG and NPAtlas for improving our ML method.^30,52^

## Conclusion

We used ML-guided genome mining in combination with bioactivity-guided fractionation to guide the isolation of two antibacterial compounds, dipyrimicins A (**1**) and B (**2**). While dipyrimicins were originally isolated from a related *Amycolatopsis* sp. K16-0194, this marks the first time 2,2’-BP family compounds and the corresponding BGC have been identified from *Amycolatopsis azurea* DSM 43854. Among the 25+ BGCs within *A. azurea* DSM 43854 predicted to encode an antimicrobial natural product, we de-orphaned a cluster with 76% activity probability as the *dip* BGC. Importantly, this represents the first example of a BGC identified using this ML method that was not part of the model’s training set. The *dip* BGC shares only 40–52% similarity to previously characterized 2,2’-BP BGCs, validating the model’s ability to predict bioactive compounds from previously uncharacterized BGCs.

To confirm the link between dipyrimicins and the *dip* BGC, we performed *in silico* sequence and structural analyses, identifying key tailoring enzymes including an O-methyltransferase and amidotransferase whose functions align with the structural modifications observed in dipyrimicins A and B. Specifically, the dipyrimicins differ in the presence of a carboxylic or amido group, and our findings suggest that C7’-amidation, catalyzed by Dip5, weakens antibacterial activity. This highlights the importance of tailoring enzymes modulating bioactivity. However, the biological advantage of C7’-amidation remains unknown, and further studies are needed to determine its potential significance in the physiology and ecology in *A. azurea* DSM 43854. Also, we expanded our ML-guided genome mining to additional 2,2’-BP BGCs and identified novel tailoring enzymes that could further contribute to structural and functional diversity within this class of natural products. Future studies can benefit from incorporating both specific tailoring genetic information and SAR trends to refine ML methods and provide insight into structural modifications that modulate bioactivity.

This study demonstrates how ML-guided approaches can be leveraged for natural product discovery. While the effort described in this paper resulted in the discovery of a known compound, it did enable the deorphanization of the compound’s BGC. We believe that this approach has broad applicability in prioritizing strains capable of producing bioactive natural products, particularly in less-studied strains where genomic information is available but their metabolites remain uncharacterized. We will continue to apply our ML genome mining pipeline with the goal of identifying novel bioactive secondary metabolites. Additionally, future studies will focus on integrating this approach with other natural product discovery strategies such as high throughput activity-screenings and heterologous expression for more targeted prioritization.

## Materials and Methods

### General experimental procedures and materials

*Amycolatopsis azurea* DSM 43854 strain used throughout this study was obtained from the German Collection of Microorganisms and Cell Cultures (DSMZ). The *Bacillus spizizenii* ATCC 6633 indicator strain was obtained from Fisher Scientific. The remaining indicator strains used for bioactivity studies, including *Escherichia coli* MG1655 ATCC 700926, *Pseudomonas aeruginosa* ATCC 27853, and *Staphylococcus aureus* ATCC 25923, were obtained from the American Type Culture Collection (ATCC). BD Difco ISP2 and ISP4 media for bacterial growth were purchased from Fisher Scientific along with the HyperSep C18 cartridges. Oxoid TSB media used for MIC assay and the HyperSIL GOLD C18 columns were acquired from Thermo Scientific. All solvents used for organic extractions, chromatography, and spectrometry experiments were HPLC and LC-MS grade materials and were purchased from Fisher Scientific and Sigma Aldrich. NMR solvents were purchased from Sigma Aldrich.

### Genome mining and BGC activity prediction

The genome sequence of *A. azurea* DSM 43854 (NCBI RefSeq assembly GCF_001995215.1) was downloaded from the NCBI database. As the machine learning algorithm by Walker and Clardy was trained on gene annotations by antiSMASH 5 and resistance gene identifier (RGI) version 5, these same versions were used for the activity prediction. ^4,8,9^ For the BGCs of interest, no notable differences were observed between the results from antiSMASH versions 5 and 7. Therefore, AntiSMASH 5 was retained for the majority of the bioinformatic analysis to ensure consistency.

The BGCs of collismycin A (Genbank accession no. HE575208.1)^22^ and caerulomycin A BGC (Genbank accession no. JF419316.2)^27^ were aligned with the putative dipyrimicin BGC using clinker through the CAGECAT webserver (https://cagecat.bioinformatics.nl/).^50^ To identify other putative producers of 2,2’-bipyridine-containing compounds, using the 2,2’-BP forming genes as a query, a combination of cblaster^53^, Enzyme Function Initiative-Enzyme Similarity Tool (EFI-EST)^53,54^, and AntiSMASH 7 (BLAST against antiSMASH-database)^55^ were used. Clinker, through the CAGECAT webserver, was used to align and confirm that each BGC, at the least, contains the conserved 2,2’-BP genes.^50^

For the structural analysis, the online version of ColabFold and its default parameters were used to generate the AlphaFold2 3D structures from the amino acid sequences of the *dip* genes.^37,56^ Structural models relevant to our analyses were identified and structurally aligned using the FoldSeek (TM-align mode) and FoldMason webserver (https://search.foldseek.com).^38,57^

### Fermentation

A frozen glycerol stock of *A. azurea* DSM 43854 was streaked onto ISP2 agar plates (4.0 g yeast extract, 10.0 g malt extract, 4.0 g glucose, 20.0 g agar) at 30°C until individual colonies were visible. A single colony was used to inoculate 5 mL of liquid ISP2 medium as seed culture and incubated at 30°C overnight (200 rpm). The seed culture was used to inoculate ISP4 agar plates (10.0g soluble starch, 1.0g K_2_HPO_4_, 1.0 MgSO_4_·7H_2_O, 0.1 mg MnCl_2_·4H_2_O, 0.1 mg ZnSO_4_·7H_2_O, 20.0 g agar) and incubated at 30°C for 14 days.

### Extraction and Isolation

After 14-day fermentations, the agar plates were frozen (-20°C) overnight before thawing for >5 hrs prior to extraction. The agar plates were then soaked in equal volumes of ethyl acetate (1L per 20 plates) and deionized water and sonicated for ∼1 hr. The liquid extract was separated from the solid agar via vacuum filtration and the ethyl acetate layer was further separated via liquid-liquid extraction. The organic fraction was concentrated *in vacuo*. The sample was redissolved in 25% ACN to obtain the crude extract for further analysis and isolation.

### Bioactivity-guided isolation

The crude extract underwent further separation through rounds of solid-phase extraction (SPE), flash chromatography, and semi-preparative HPLC. Dipyrimicin A and B were identified as bioactive compounds of interest via bioactivity-guided fractionation in which each chromatography fractions obtained were concentrated *in vacuo* then subjected to a bioassay initially against *Bacillus spizizenii* ATCC 6633. Those showing activity (any observed zone of inhibition greater than the 25% ACN negative control) were further fractionated to obtain individual bioactive metabolites. SPEs were performed using HyperSep C18 Cartridges (5000 mg bed weight). The cartridge was washed with 2-3 column volumes (CV) of 100% Methanol and equilibrated with 2 CVs of water before loading the crude sample. Analytes were then eluted from the column using 25%, 50%, 75%, 100% ACN. Fractions collected at 50% ACN displayed large zone of inhibition and were subjected to flash chromatography. Büchi Pure C-850 FlashPrep system equipped with UV and ELSD detectors and FlashPure Ecoflex C18 cartridge (50 μm, spherical, 4 g) were used in linear gradient from 0 to 100% acetonitrile (ACN) and decreasing volumes of H_2_O in 10 min. Bioactivity was observed on fractions eluting at around 3 min and were advanced for further HPLC purifications. These were carried out on a Thermo Fisher Vanquish HPLC system equipped with a photodiode array detector and an automated fraction collector using a Hypersil GOLD C18 column (3 um, 4.6 x 100 mm) with a linear gradient of 0 to 100% ACN/H_2_O (95/5) containing 0.1% (v/v) formic acid (FA) in 10 min. Individual peaks corresponding to our active compounds of interest were collected at around 5.5 to 6.5 min.

### Characterization

UV-Vis spectra were recorded from 190 - 500 nm on a Thermo Fisher Vanquish HPLC system equipped with a photodiode array detector. An analytical Hypersil GOLD C18 column (3 μm, 4.6 x 100 mm) was used to resolve samples at a flow rate of 1 ml/min with a linear gradient from 0 to 100% ACN/H_2_O (95/5) containing 0.1% (v/v) formic acid (FA) in 10 min. ESI-HRMS were measured on a LTQ**-**Orbitrap 3 XL at the Vanderbilt University Mass Spectrometry Core Lab. Data was acquired in positive ion mode between 100-2000 *m/z*. NMR spectra, including ^1^H, ^1^H-^1^H COSY, ^1^H-^13^C HSQC, and ^1^H-^13^C HMBC experiments, were recorded in CD_3_OD on a 600 MHz NMR spectrometer (Bruker) at the Vanderbilt University Small Molecule NMR Facility Core.

### Antimicrobial assays

The minimum inhibitory concentration (MIC) was defined as the lowest concentration of dipyrimicin inhibiting visible growth after incubation. MICs to the indicator strains *Bacillus spizizenii* ATCC 6633, *Escherichia coli* MG1655 ATCC 700926, *Pseudomonas aeruginosa* ATCC 27853, and *Staphylococcus aureus* ATCC 25923 were determined following the guidelines of the Clinical Laboratory Standards Institute (CLSI) with minor modifications. The indicator strains were grown in Tryptic Soy Broth (TSB) overnight at 30°C with shaking. Dipyrimicin A and B were 10-fold serially diluted from 256 μg/mL to 0.5 μg/mL in a 96-well plate using TSB with a total volume of 50uL per well. The inoculum was prepared according to Kadeřábková et al.^58^ and 50uL per well were added which diluted the dipyrimicins to reach the final test concentrations of 128 μg/mL to 0.25 μg/mL. The plates were incubated at 30°C for 24h and the absorbance at OD_600_ was monitored using Varioskan LUX Multimode Microplate Reader. Tetracycline was used as a positive control. Each sample concentration was run in triplicate.

## Acknowledgements

Research reported in this publication was supported by the National Institute of General Medical Sciences under award number R35GM146987. The content is solely the responsibility of the authors and does not necessarily represent the official views of the National Institutes of Health. We would also like to acknowledge the Littlejohn Fellowship for supporting Isra Shuster’s participation in the Vanderbilt Undergraduate Summer Research Program, Vanderbilt’s ACCRE and Center for Structural Biology for computational resources, and Dr. Donald Stec for advice on NMR experiments.

