## Supplementary information for "AI-assisted isolation of bioactive Dipyrimicins from *Amycolatopsis azurea* and identification of its corresponding *dip* biosynthetic gene cluster"

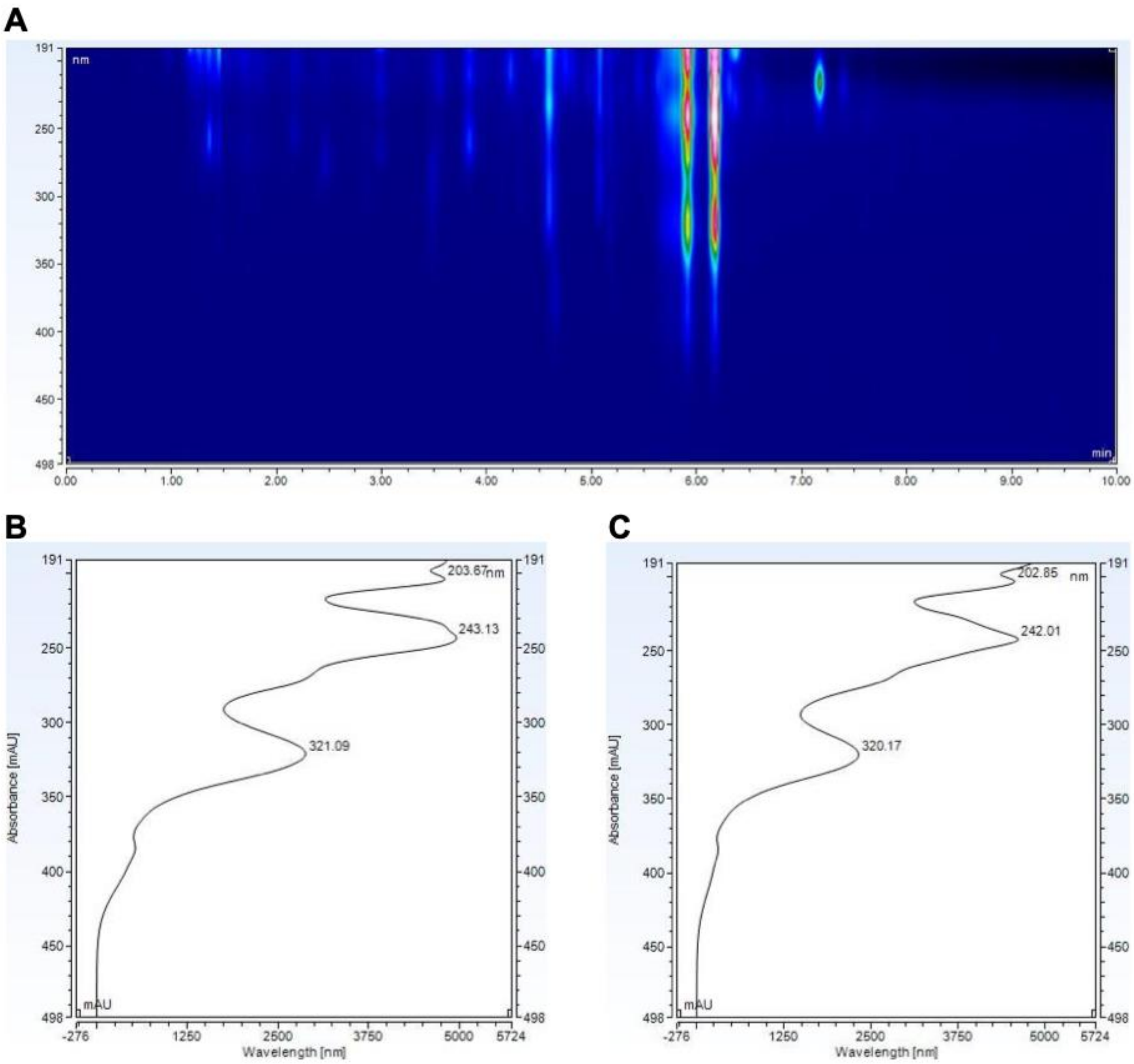


**Figure S1. (A)** HPLC chromatogram and UV-Vis spectrum of *A. azurea* DSM 43854 culture extract showing typical UV-Vis spectrum for 2,2'-bipyridines (**B**) Dipyrimicin A (compound **1,** λ = 204nm, 243nm, 268nm, and 321nm, retention time (RT) at ~5.8 min) and (**C**) Dipyrimicin B (compound **2,** λ = 203nm, 242nm, 268nm, and 320nm, RT at ~6.2 min).

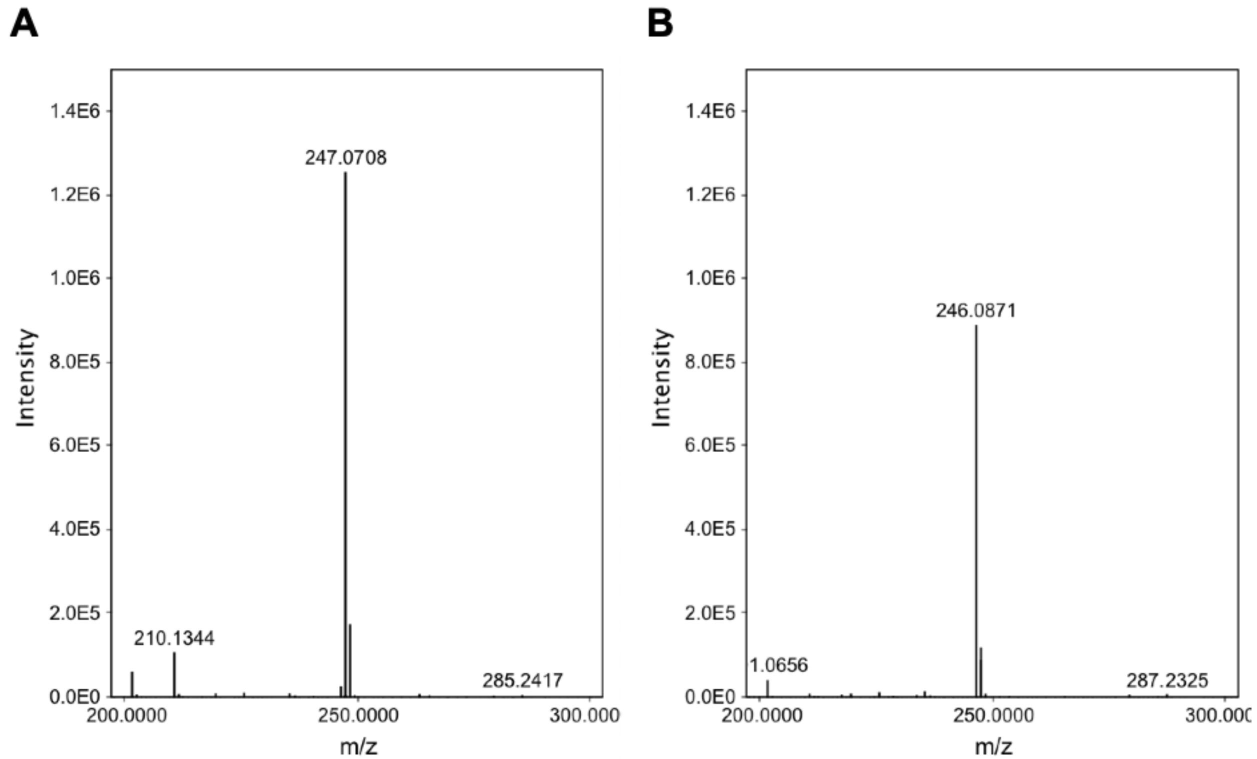


**Figure S2.** HR-ESI-MS spectrum of **(A)** Dipyrimicin A (compound **1,** m/z $[M+H]^+$ obsvd. 247.0708, calcd. for $C_{12}H_{10}N_2O_4$, 247.0719) and (**B**) Dipyrimicin B (compound **2,** m/z $[M+H]^+$ obsvd. 246.0871, calcd. for $C_{12}H_{11}N_3O_3$, 246.0879)

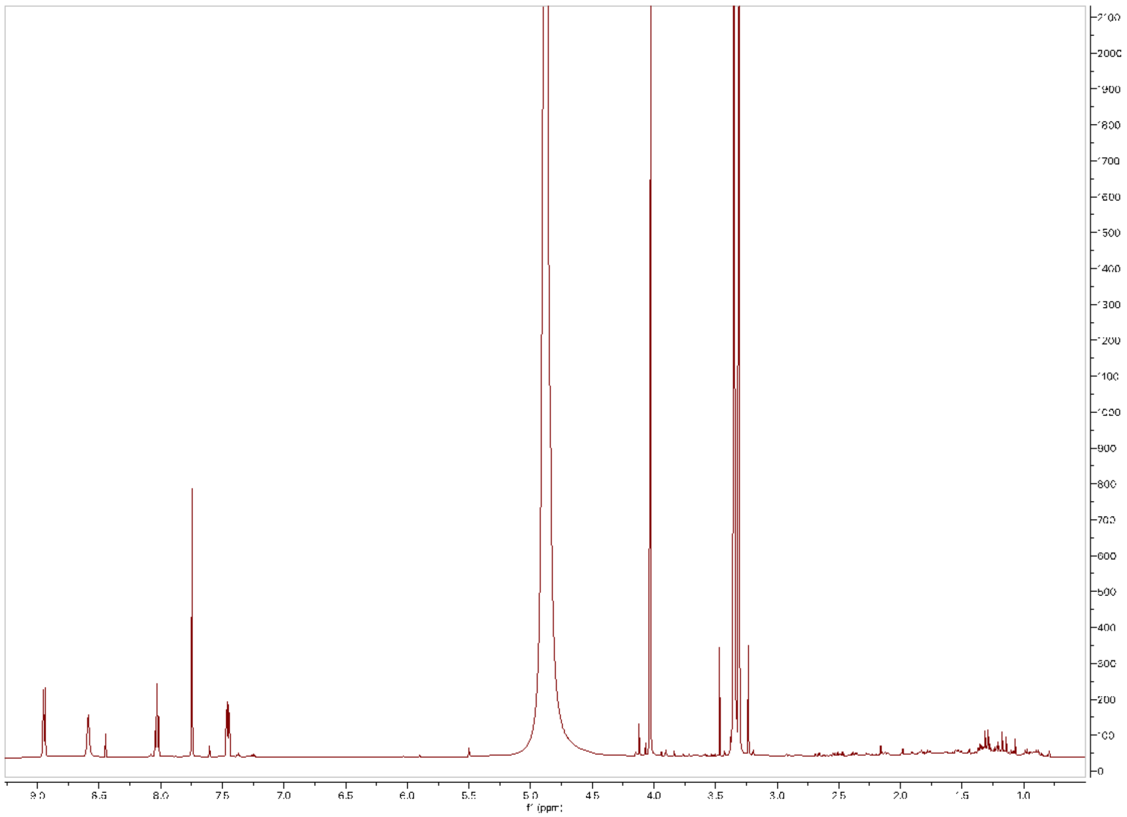

**Figure S3.** $^{1}$H NMR (600 MHz) spectrum of Dipyrimicin A (**1**) in $CD_3OD$.

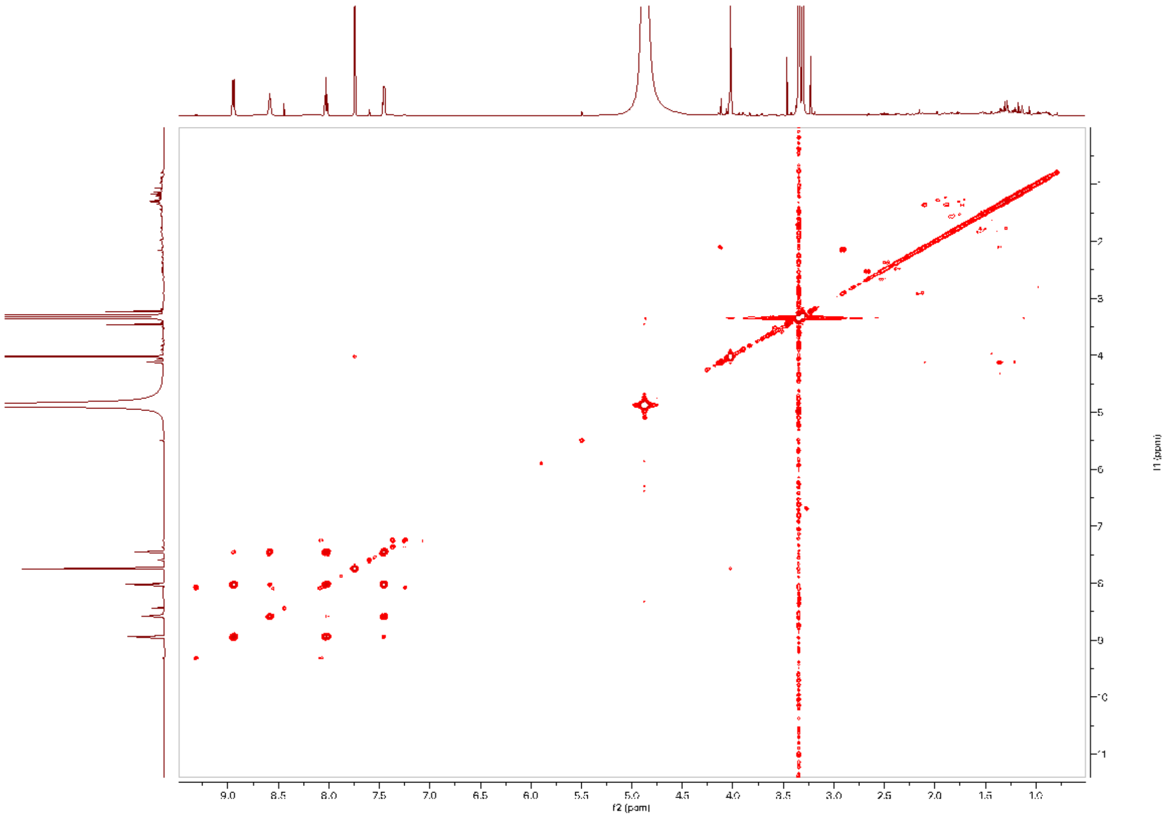


**Figure S4.** $^{1}H$-$^{1}H$ COSY NMR (600 MHz) spectrum of Dipyrimicin A (**1**) in $CD_3OD$.

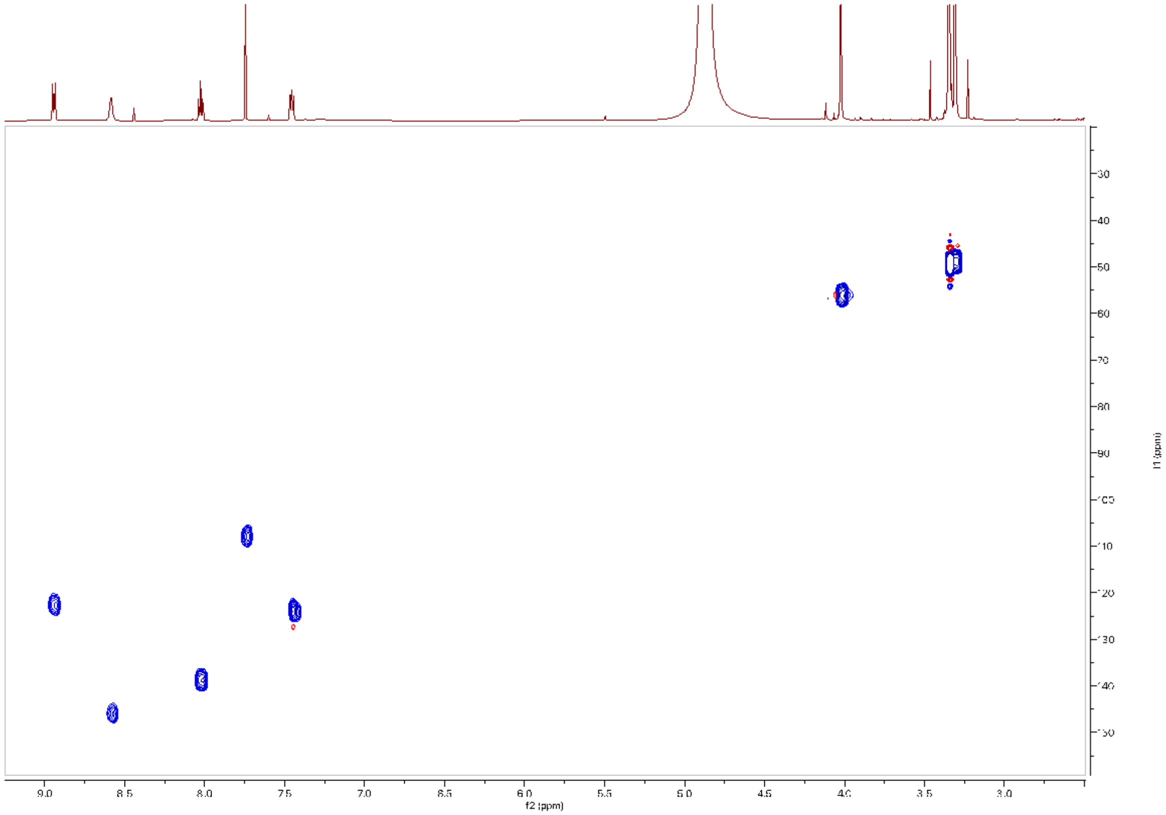

**Figure S5.** $^{1}H$-$^{13}C$ HSQC NMR (600 MHz) spectrum of Dipyrimicin A (**1**) in $CD_3OD$.

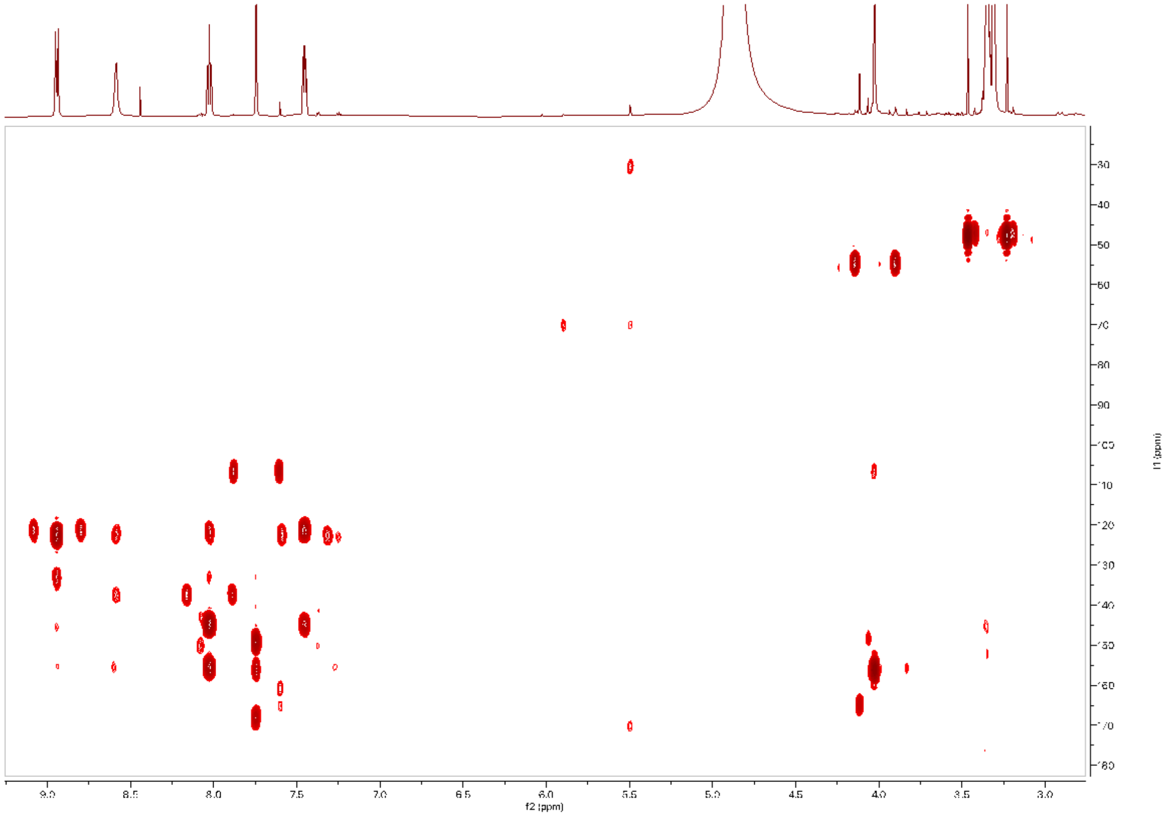

**Figure S6.** $^{1}H$-$^{13}C$ HMBC NMR (600 MHz) spectrum of Dipyrimicin A (**1**) in $CD_3OD$.

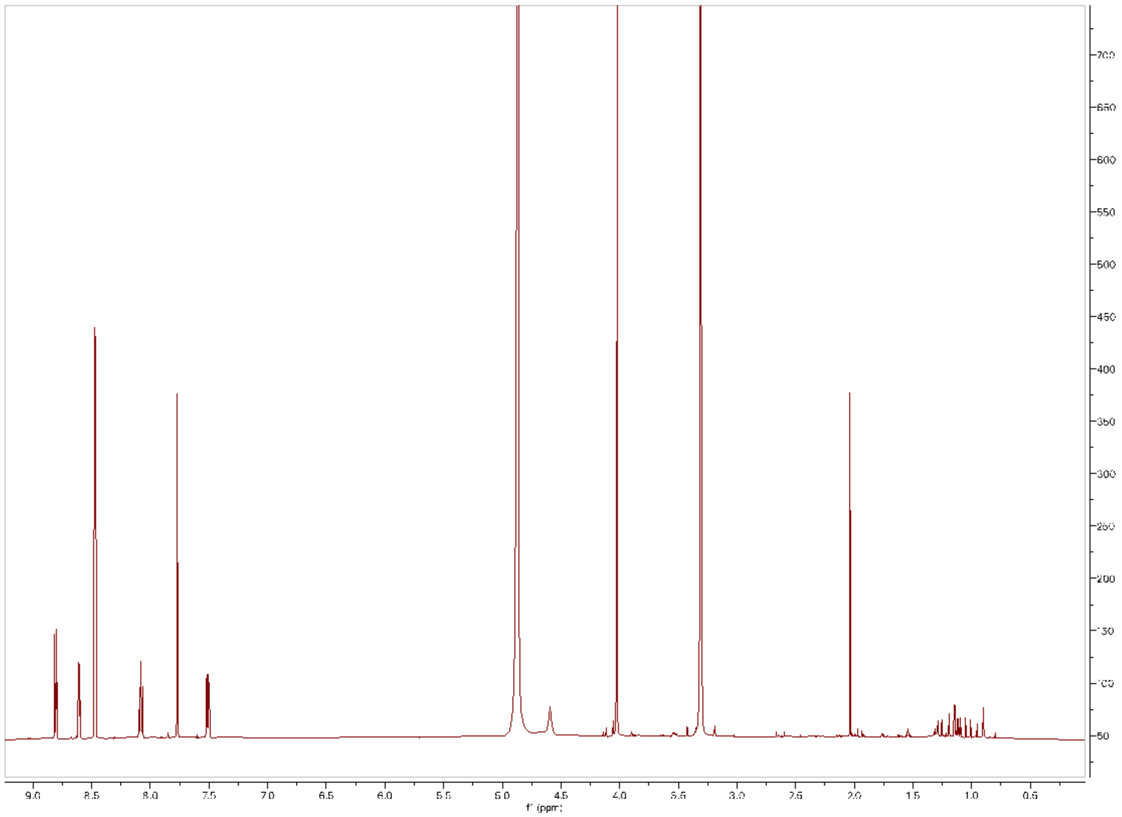


**Figure S7.** $^{1}H$ NMR (600 MHz) spectrum of Dipyrimicin B (**2**) in $CD_3OD$.

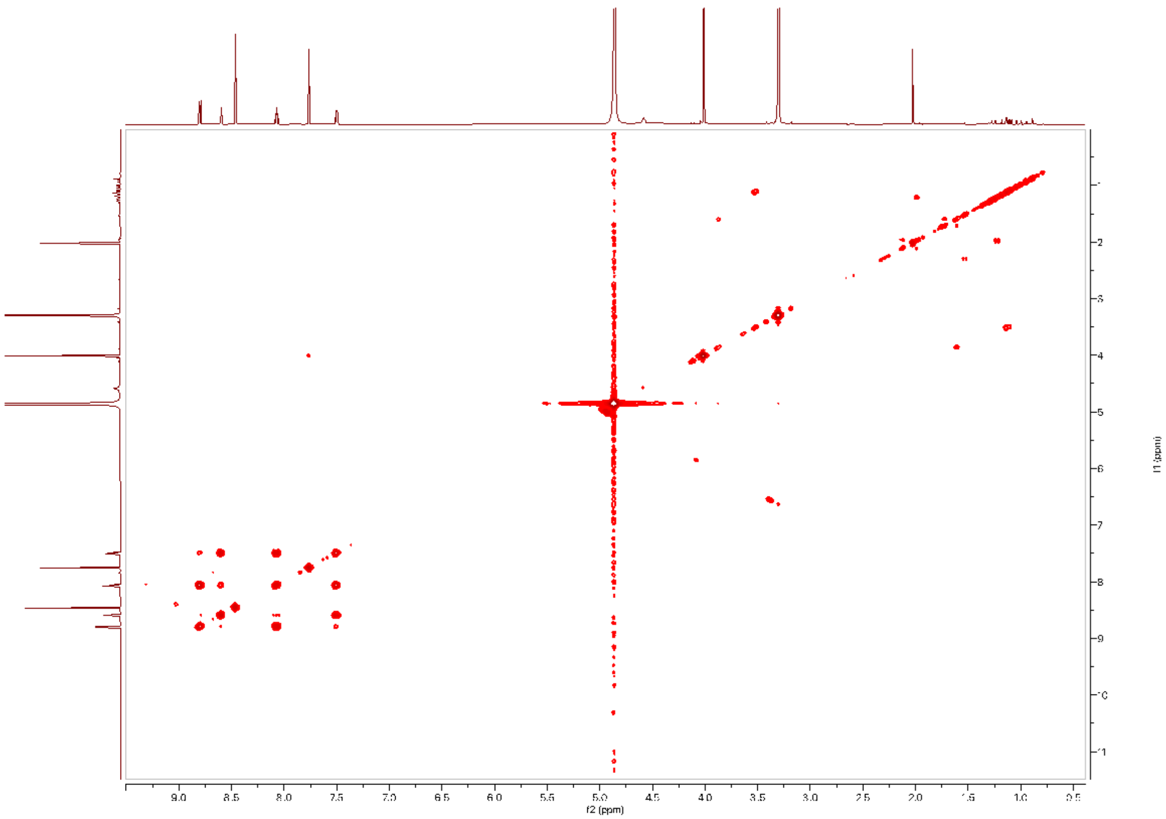


**Figure S8.** $^{1}H$-$^{1}H$ COSY NMR (600 MHz) spectrum of Dipyrimicin B (**2**) in $CD_3OD$.

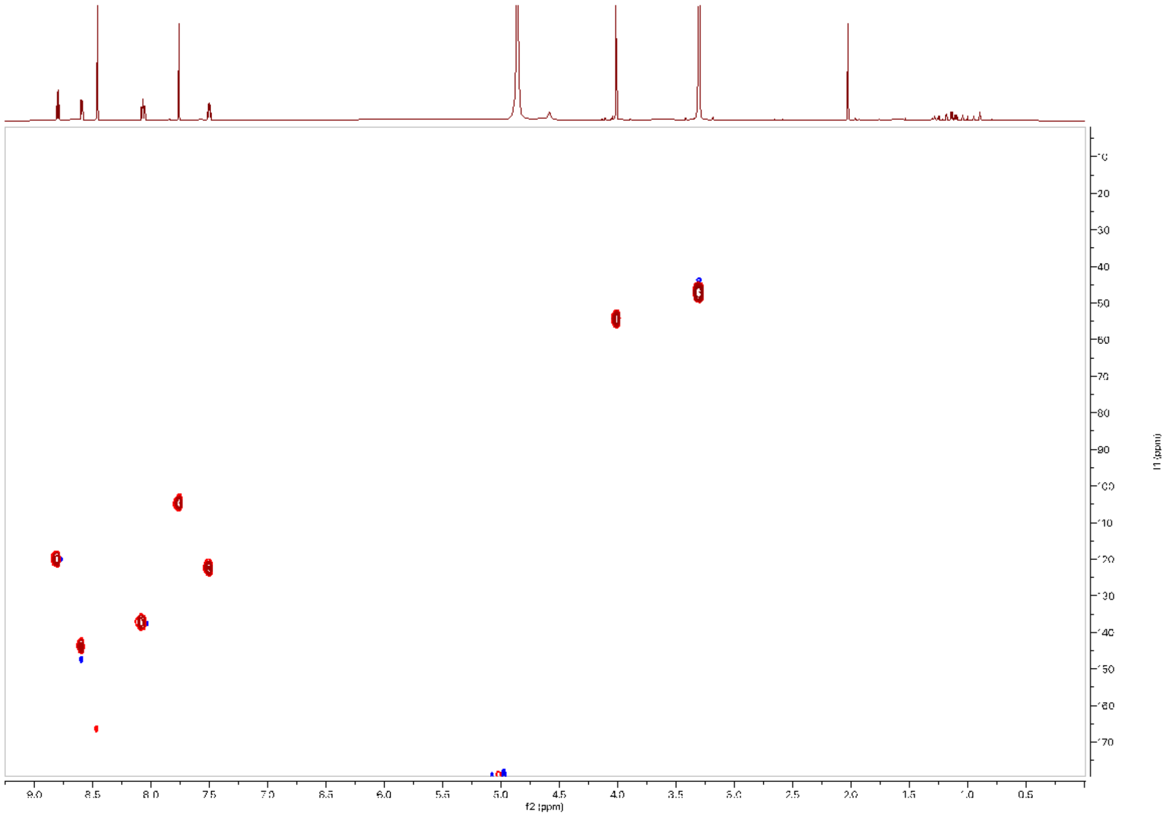

**Figure S9.** $^{1}$H-$^{13}$C HSQC NMR (600 MHz) spectrum of Dipyrimicin B (**2**) in $CD_3OD$.

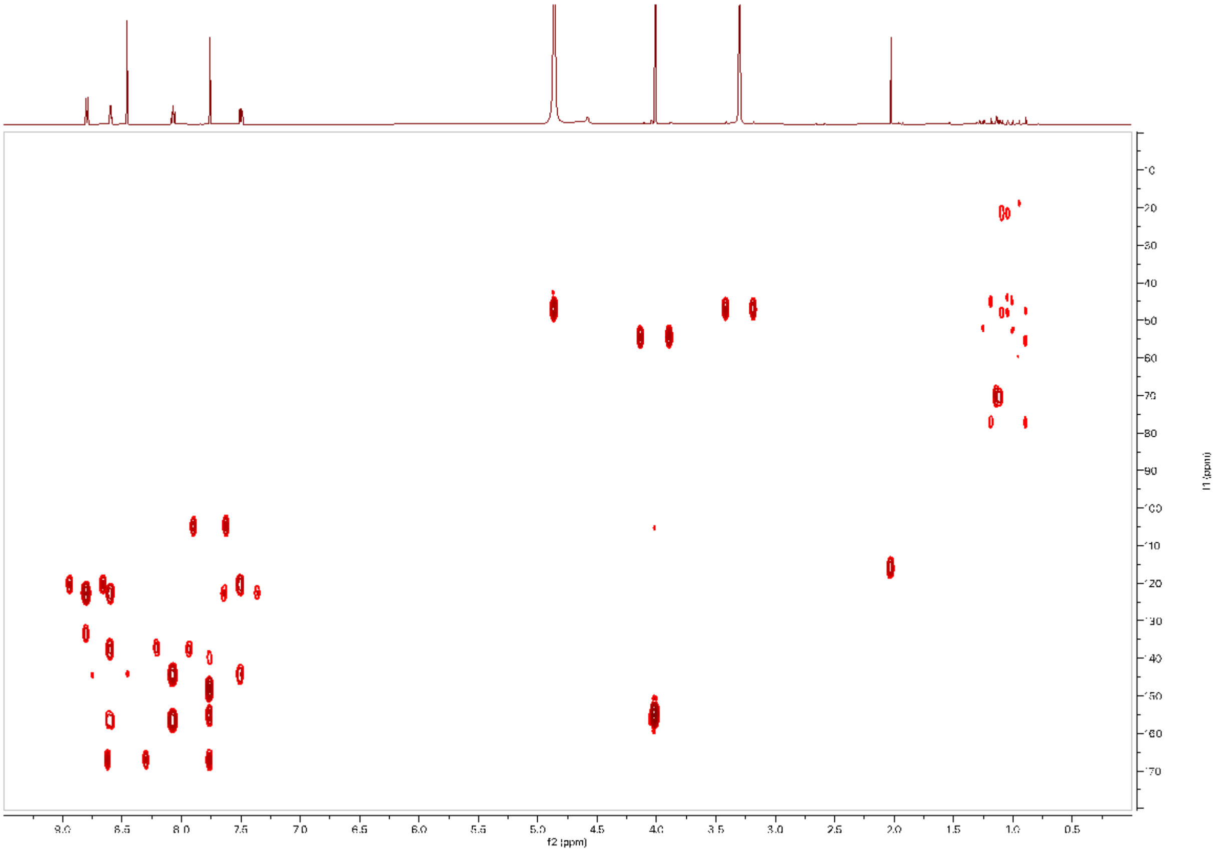


**Figure S10.** $^{1}$H-$^{13}$C HMBC NMR (600 MHz) spectrum of Dipyrimicin B (**2**) in $CD_3OD$.

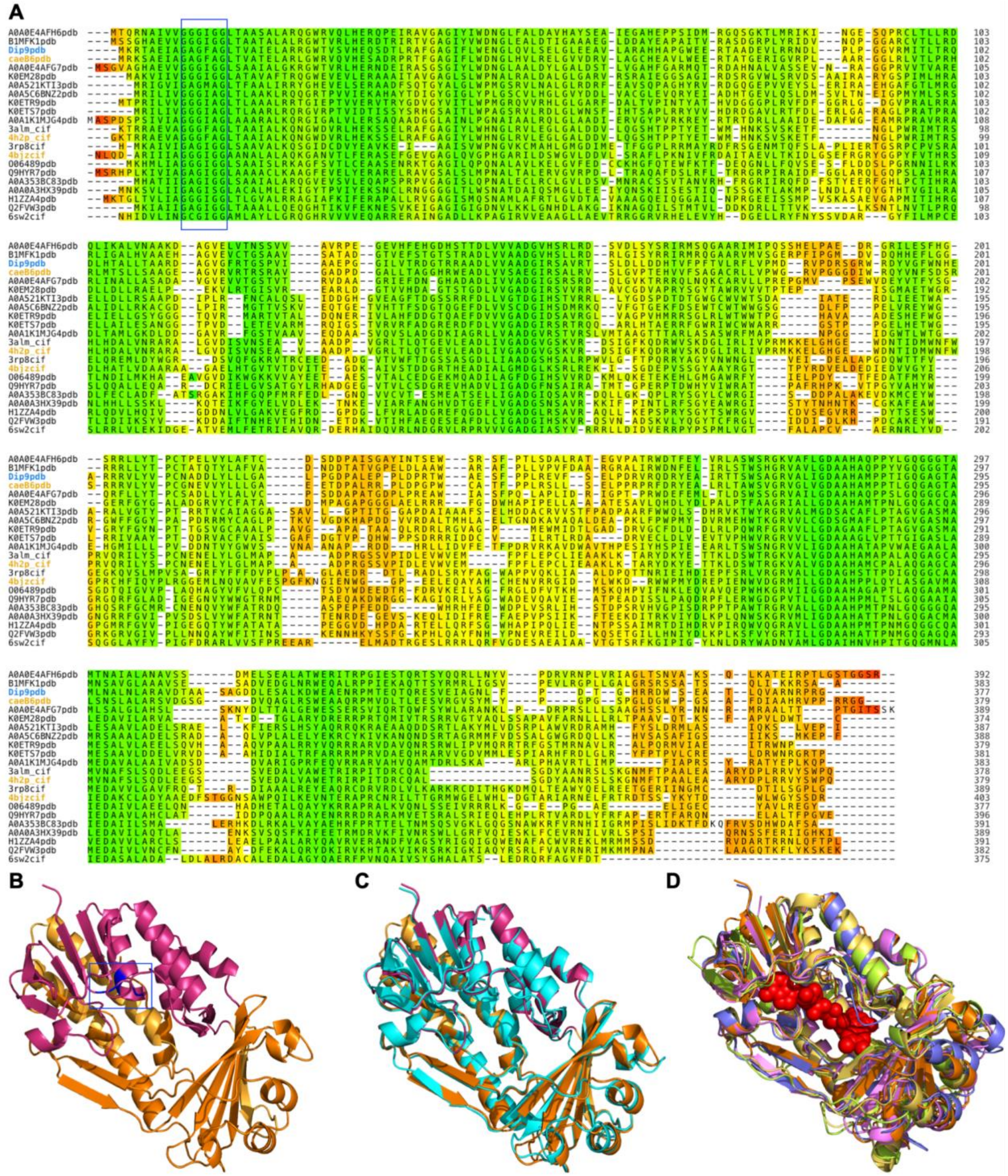

**Figure S11.** Multiple structural alignment (MSTA) of Dip9 and related hydroxylases. A) FoldMason MSTA of the Dip9 structure along with the top 5 FoldSeek hits from different databases (**Table S4**). Residues are shaded by the LDDT score of their alignment column (color scheme: red 0%, green 100%). Highlighted in the blue box is the conserved “GxGxxG” motif found among flavin-containing oxygenases.[1] B) The AlphaFold-predicted structure of Dip9. Highlighted in pink and dark orange are the putative FAD-binding and substrate-binding domains, respectively. The glycine-rich motif is depicted in blue within the box. C) Superimposition of the predicted structures of Dip9 and CaeB6 (cyan). D) Superimposition of Dip9 and the top FoldSeek hits from different databases (**Table S4**) and the representative FAD is depicted in red spheres (general location of FAD in PDBs 4H2P and 4BJZ), highlighting that the related hydroxylases have similar sites for FAD- and substrate-binding.

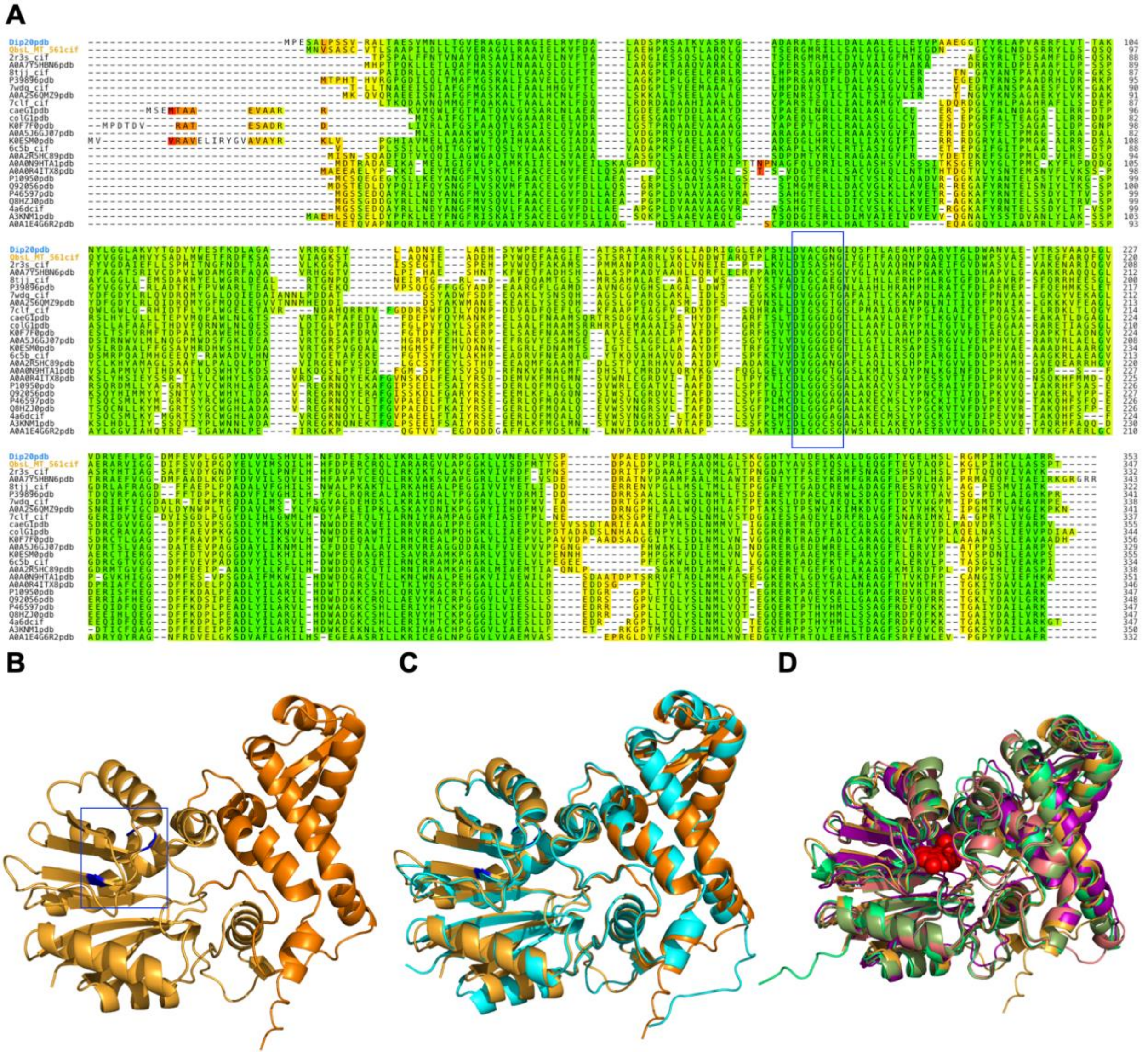


**Figure S12.** Multiple structural alignment (MSTA) of Dip20 and related methyltransferases. A) FoldMason MSTA of the Dip20 structure along with the top 5 FoldSeek hits from different databases (**Table S5**). Residues are shaded by the LDDT score of their alignment column (color scheme: red 0%, green 100%). Highlighted in the blue box is the "DxGxGxG" or "GxG" fingerprint, a SAM-binding motif found among class I methyltransferases. B) The AlphaFold-predicted structure of Dip20. The C-terminal domain (light orange) contains the pocket for SAM- and substrate-binding while the N-terminal domain is for dimerization.The glycine-rich motif is depicted in blue within the box. C) Superimposition of the predicted structures of Dip20 and the C-terminal domain of QbsL (cyan). D) Superimposition of Dip20 and the top FoldSeek hits from different databases (**Table S5**). The representative SAM molecule is depicted in red spheres (from 2R3S PDB).

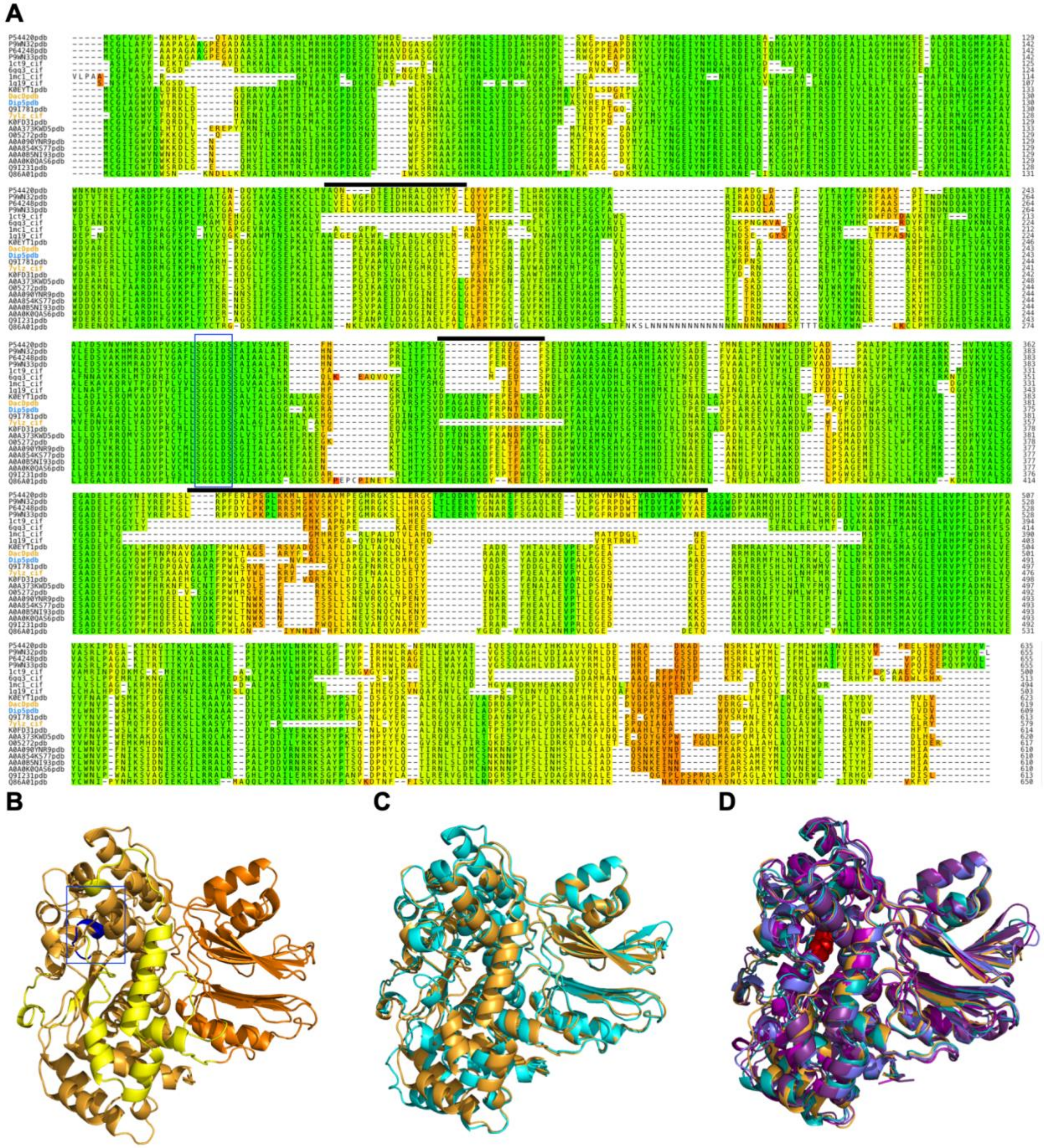

**Figure S13.** Multiple structural alignment (MSTA) of Dip5 and related amidotransferases. A) FoldMason MSTA of the Dip5 structure along with the top 5 FoldSeek hits from different databases (**Table S6**). Residues are shaded by the LDDT score of their alignment column (color scheme: red 0%, green 100%). Highlighted in the blue box is the conserved "SGGLDS" motif among amidotransferases. Dip5 and several other related proteins reveal inserted regions (black bars) that are absent when compared to the glutamine-dependent asparagine synthetase, AsnB, from *Escherichia coli* (PDB 1CT9). B) The AlphaFold-predicted structure of Dip5. Highlighted in light and dark orange are the C-terminal synthetase and N-terminal glutaminase domains, respectively. Residues in yellow are the inserted regions not found in AsnB. The conserved motif involved in ATP-binding is depicted in blue within the box. C) Superimposition of the predicted structures of Dip5 and DacD. D) Superimposition of Dip5 and the top FoldSeek hits from different databases (**Table S6**). The representative ATP molecule is depicted in red spheres (from 1CT9 PDB).

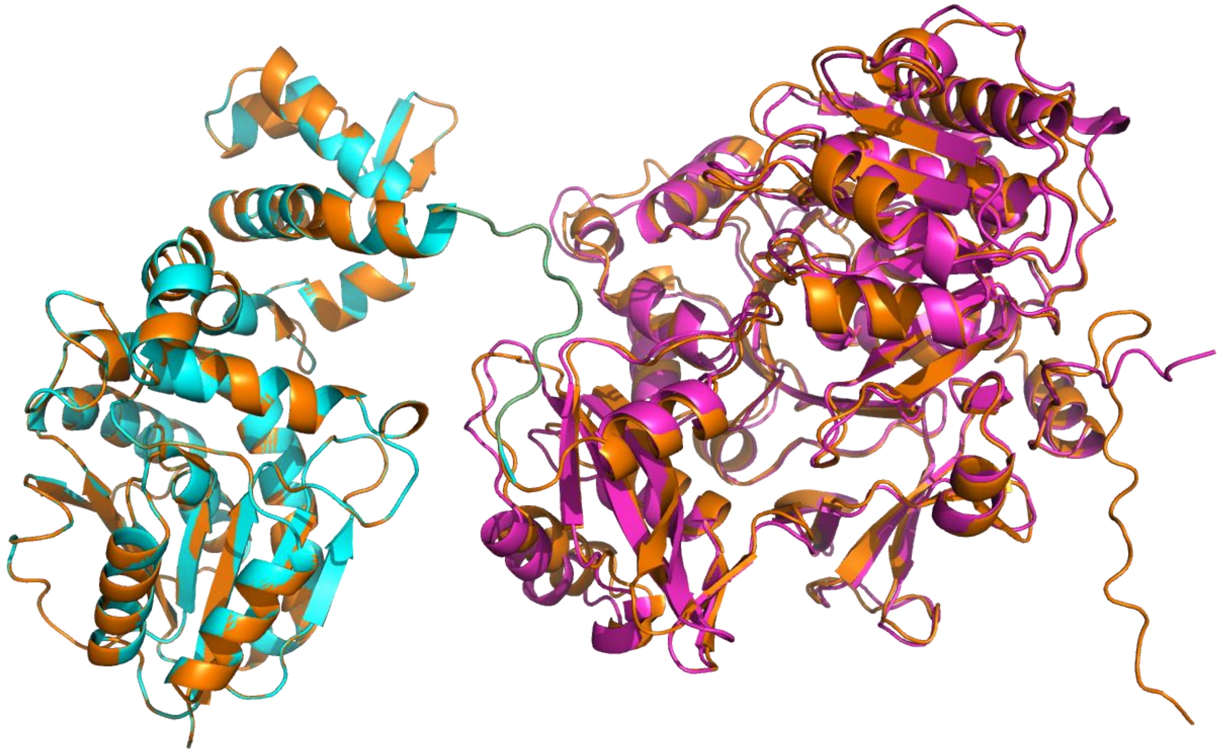

**Figure S14.** Superimposition of the AlphaFold-predicted structures of Dip21 (magenta) and Dip20 (cyan) to the N-terminal synthetase and C-terminal methylase domains of QbsL (orange), respectively.

**Table S1.** The biosynthetic gene clusters (BGCs) of secondary metabolites in *Amycolatopsis azurea* DSM 43854 analyzed from antiSMASH 5.

| Cluster | Type | From | To | Most similar known BGC (% similarity) |
|---|---|---|---|---|
| 1.1 | CDPS | 137,563 | 158,264 | |
| 1.2 | terpene | 210,895 | 233,051 | geosmin (100%) |
| 1.3 | ectoine | 256,222 | 266,611 | ectoine (100%) |
| 2.1 | T1PKS | 318 | 68,485 | butyrolactol A (66%) |
| 2.2 | hglE-KS, T1PKS | 108,031 | 158,424 | rimosamide (14%) |
| 2.3 | lassopeptide | 268,408 | 290,889 | ansacarbamitocin A (12%) |
| 2.4 | NRPS, amglyccycl | 324,333 | 378,452 | acarbose (14%) |
| 2.5 | NRPS, T1PKS | 378,800 | 487,216 | caerulomycin A (52%) |
| 2.6 | NRPS | 548,461 | 609,364 | |

| | | | | |
|---|---|---|---|---|
| 2.7 | T1PKS | 610,946 | 657,329 | kanamycin (1%) |
| 5.1 | lanthipeptide | 90,029 | 112,572 | Ery-9 / Ery-6 / Ery-8 / Ery-7 / Ery-5 / Ery-4 / Ery-3 (75%) |
| 6.1 | terpene | 166,574 | 187,503 | isorenieratene (42%) |
| 7.1 | NRPS-like | 10,140 | 54,042 | ansacarbamitocin A (4%) |
| 10.1 | NRPS | 16,609 | 84,811 | mirubactin (78%) |
| 11.1 | terpene | 23,959 | 46,151 | enduracidin (10%) |
| 11.2 | betalactone | 66,497 | 91,707 | |
| 12.1 | arylpolyene | 78,681 | 119,832 | kinamycin (5%) |
| 15.1 | NRPS | 116,412 | 178,671 | albachelin (100%) |
| 16.1 | bacteriocin | 162,646 | 173,458 | |
| 16.2 | terpene | 364,878 | 386,005 | vazabitide A (4%) |
| 16.3 | T1PKS | 432,515 | 469,723 | funisamine (22%) |
| 20.1 | other, arylpolyene | 21,861 | 73,484 | kedarcidin (19%) |
| 22.1 | NRPS-like | 181,449 | 225,240 | meilingmycin (3%) |
| 23.1 | NRPS, CDPS, T1PKS, T3PKS, terpene | 1,589 | 122,641 | diazepinomicin (17%) |
| 24.1 | terpene | 65,035 | 86,174 | 2-methylisoborneol (100%) |
| 24.2 | NRPS-like, lanthipeptide | 108,423 | 151,788 | |
| 24.3 | NRPS | 214,282 | 273,246 | capreomycin IA / capreomycin IB / capreomycin IIA / capreomycin IIB (9%) |
| 25.1 | T3PKS, NRPS-like, NRPS | 124,067 | 226,706 | keratinimicin A / keratinimicin B / keratinimicin C / keratinimicin D (92%) |
| 25.2 | T1PKS, NRPS-like | 233,345 | 279,671 | polyoxypeptin (10%) |
| 27.1 | lanthipeptide | 16,974 | 39,844 | |
| 28.1 | T1PKS | 3,842 | 49,676 | amycolamycin A / amycolamycin B (31%) |
| 28.2 | T2PKS | 134,047 | 206,532 | actinorhodin (59%) |
| 29.1 | hglE-KS, T1PKS | 14,179 | 64,527 | rifamorpholine A / rifamorpholine B / rifamorpholine C / rifamorpholine D / rifamorpholine E (9%) |
| 33.1 | T1PKS, NRPS | 1 | 46,481 | chlortetracycline (5%) |
| 45.1 | T1PKS | 1 | 26,493 | |

**Table S2.** Machine-learning predicted probability scores for various bioactivities of the AntiSMASH 5.0 annotated BGCs from *A. azurea DSM 43854*. The scores were obtained using three classifiers [extra trees, logistic regression, and support vector machine (SVM)] and were averaged for each activity classification.

| | Antibacterial | | | | Anti-gram positive | | | | Anti-gram negative | | | | antifungal/antitumor/cytotoxic | | | | Antifungal | | | | Antitumor/cytotoxic | | | |
|---|---|---|---|---|---|---|---|---|---|---|---|---|---|---|---|---|---|---|---|---|---|---|---|---|
| **Cluster** | **Tree** | **Log** | **SVM** | **Avg** | **Tree** | **Log** | **SVM** | **Avg** | **Tree** | **Log** | **SVM** | **Avg** | **Tree** | **Log** | **SVM** | **Avg** | **Tree** | **Log** | **SVM** | **Avg** | **Tree** | **Log** | **SVM** | **Avg** |
| 1.1 | 0.68 | 0.50 | 0.56 | 0.58 | 0.56 | 0.27 | 0.28 | 0.37 | 0.24 | 0.05 | 0.04 | 0.11 | 0.36 | 0.38 | 0.33 | 0.36 | 0.19 | 0.13 | 0.05 | 0.12 | 0.26 | 0.28 | 0.30 | 0.28 |
| 1.2 | 0.35 | 0.37 | 0.60 | 0.44 | 0.48 | 0.28 | 0.35 | 0.37 | 0.12 | 0.15 | 0.19 | 0.15 | 0.28 | 0.19 | 0.18 | 0.22 | 0.20 | 0.23 | 0.18 | 0.20 | 0.10 | 0.14 | 0.12 | 0.12 |
| 1.3 | 0.46 | 0.35 | 0.17 | 0.33 | 0.40 | 0.15 | 0.17 | 0.24 | 0.00 | 0.09 | 0.09 | 0.06 | 0.20 | 0.23 | 0.22 | 0.22 | 0.07 | 0.16 | 0.09 | 0.11 | 0.03 | 0.06 | 0.05 | 0.05 |
| 2.1 | 0.36 | 0.28 | 0.42 | 0.35 | 0.44 | 0.11 | 0.20 | 0.25 | 0.12 | 0.15 | 0.17 | 0.15 | 0.80 | 0.89 | 0.92 | 0.87 | 0.42 | 0.54 | 0.34 | 0.43 | 0.44 | 0.25 | 0.44 | 0.38 |
| 2.2 | 0.66 | 0.53 | 0.62 | 0.60 | 0.70 | 0.46 | 0.51 | 0.56 | 0.60 | 0.35 | 0.23 | 0.39 | 0.80 | 0.69 | 0.75 | 0.75 | 0.46 | 0.36 | 0.30 | 0.37 | 0.38 | 0.12 | 0.23 | 0.24 |
| 2.3 | 0.90 | 0.86 | 0.98 | 0.91 | 0.78 | 0.98 | 0.92 | 0.89 | 0.48 | 0.36 | 0.27 | 0.37 | 0.44 | 0.10 | 0.10 | 0.21 | 0.16 | 0.12 | 0.26 | 0.18 | 0.21 | 0.04 | 0.20 | 0.15 |
| 2.4 | 0.72 | 0.75 | 0.72 | 0.73 | 0.62 | 0.39 | 0.51 | 0.51 | 0.52 | 0.92 | 0.43 | 0.62 | 0.72 | 0.96 | 0.93 | 0.87 | 0.51 | 0.45 | 0.25 | 0.40 | 0.64 | 0.04 | 0.35 | 0.35 |
| 2.5 | 0.60 | 0.99 | 0.71 | 0.76 | 0.60 | 1.00 | 0.95 | 0.85 | 0.40 | 0.91 | 0.42 | 0.58 | 0.72 | 0.92 | 0.93 | 0.86 | 0.62 | 0.27 | 0.17 | 0.35 | 0.50 | 0.07 | 0.35 | 0.31 |
| 2.6 | 0.64 | 0.52 | 0.59 | 0.58 | 0.48 | 0.35 | 0.32 | 0.38 | 0.20 | 0.10 | 0.11 | 0.14 | 0.40 | 0.82 | 0.65 | 0.62 | 0.34 | 0.19 | 0.17 | 0.23 | 0.21 | 0.60 | 0.32 | 0.38 |
| 2.7 | 0.66 | 0.53 | 0.62 | 0.60 | 0.64 | 0.26 | 0.41 | 0.44 | 0.24 | 0.29 | 0.35 | 0.29 | 0.45 | 0.03 | 0.06 | 0.18 | 0.37 | 0.18 | 0.16 | 0.24 | 0.40 | 0.04 | 0.31 | 0.25 |
| 5 | 0.58 | 0.69 | 0.68 | 0.65 | 0.62 | 0.73 | 0.63 | 0.66 | 0.32 | 0.52 | 0.30 | 0.38 | 0.20 | 0.04 | 0.07 | 0.10 | 0.28 | 0.12 | 0.19 | 0.20 | 0.11 | 0.01 | 0.08 | 0.06 |
| 6 | 0.20 | 0.23 | 0.09 | 0.17 | 0.20 | 0.04 | 0.10 | 0.11 | 0.28 | 0.10 | 0.17 | 0.18 | 0.24 | 0.14 | 0.28 | 0.22 | 0.18 | 0.16 | 0.28 | 0.21 | 0.13 | 0.04 | 0.06 | 0.08 |
| 7 | 0.52 | 0.56 | 0.71 | 0.59 | 0.80 | 0.74 | 0.79 | 0.78 | 0.32 | 0.07 | 0.24 | 0.21 | 0.36 | 0.12 | 0.11 | 0.20 | 0.32 | 0.11 | 0.17 | 0.20 | 0.14 | 0.15 | 0.32 | 0.21 |
| 10 | 0.46 | 0.27 | 0.66 | 0.46 | 0.38 | 0.00 | 0.05 | 0.14 | 0.36 | 0.00 | 0.03 | 0.13 | 0.48 | 0.01 | 0.02 | 0.17 | 0.28 | 0.05 | 0.16 | 0.16 | 0.45 | 0.00 | 0.33 | 0.26 |
| 11.1 | 0.78 | 0.78 | 0.70 | 0.75 | 0.80 | 0.93 | 0.79 | 0.84 | 0.16 | 0.07 | 0.09 | 0.11 | 0.40 | 0.13 | 0.17 | 0.23 | 0.22 | 0.13 | 0.17 | 0.17 | 0.32 | 0.11 | 0.21 | 0.22 |
| 11.2 | 0.70 | 0.54 | 0.82 | 0.69 | 0.54 | 0.84 | 0.80 | 0.73 | 0.32 | 0.09 | 0.10 | 0.17 | 0.48 | 0.44 | 0.37 | 0.43 | 0.10 | 0.15 | 0.20 | 0.15 | 0.30 | 0.69 | 0.42 | 0.47 |
| 12 | 0.84 | 0.78 | 0.71 | 0.77 | 0.74 | 0.89 | 0.88 | 0.84 | 0.32 | 0.13 | 0.33 | 0.26 | 0.36 | 0.00 | 0.06 | 0.14 | 0.10 | 0.04 | 0.13 | 0.09 | 0.29 | 0.05 | 0.29 | 0.21 |
| 15 | 0.26 | 0.43 | 0.57 | 0.42 | 0.36 | 0.36 | 0.42 | 0.38 | 0.28 | 0.22 | 0.34 | 0.28 | 0.36 | 0.14 | 0.13 | 0.21 | 0.22 | 0.06 | 0.16 | 0.15 | 0.15 | 0.03 | 0.29 | 0.15 |
| 16.1 | 0.52 | 0.39 | 0.43 | 0.45 | 0.24 | 0.31 | 0.28 | 0.28 | 0.00 | 0.08 | 0.09 | 0.05 | 0.00 | 0.21 | 0.22 | 0.14 | 0.01 | 0.17 | 0.18 | 0.12 | 0.02 | 0.14 | 0.06 | 0.07 |
| 16.2 | 0.71 | 0.50 | 0.40 | 0.54 | 0.52 | 0.34 | 0.36 | 0.41 | 0.20 | 0.15 | 0.19 | 0.18 | 0.12 | 0.14 | 0.10 | 0.12 | 0.11 | 0.06 | 0.06 | 0.08 | 0.08 | 0.20 | 0.08 | 0.12 |
| 16.3 | 0.62 | 0.48 | 0.40 | 0.50 | 0.60 | 0.50 | 0.56 | 0.55 | 0.04 | 0.11 | 0.17 | 0.11 | 0.72 | 0.32 | 0.24 | 0.42 | 0.34 | 0.32 | 0.21 | 0.29 | 0.45 | 0.07 | 0.13 | 0.22 |
| 20 | 0.54 | 0.49 | 0.69 | 0.57 | 0.76 | 0.26 | 0.37 | 0.47 | 0.08 | 0.00 | 0.06 | 0.05 | 0.68 | 0.26 | 0.36 | 0.44 | 0.30 | 0.18 | 0.19 | 0.22 | 0.59 | 0.22 | 0.38 | 0.40 |
| 22 | 0.82 | 0.87 | 0.70 | 0.80 | 0.78 | 0.97 | 0.87 | 0.87 | 0.44 | 0.75 | 0.45 | 0.55 | 0.52 | 0.35 | 0.47 | 0.45 | 0.24 | 0.12 | 0.18 | 0.18 | 0.44 | 0.04 | 0.30 | 0.26 |
| 23 | 0.50 | 0.60 | 0.70 | 0.60 | 0.58 | 0.39 | 0.69 | 0.55 | 0.56 | 0.34 | 0.34 | 0.42 | 0.60 | 0.96 | 0.98 | 0.85 | 0.46 | 0.47 | 0.17 | 0.37 | 0.57 | 0.69 | 0.35 | 0.54 |
| 24.1 | 0.52 | 0.45 | 0.15 | 0.37 | 0.62 | 0.31 | 0.32 | 0.42 | 0.16 | 0.10 | 0.09 | 0.12 | 0.24 | 0.31 | 0.29 | 0.28 | 0.15 | 0.19 | 0.09 | 0.14 | 0.14 | 0.27 | 0.17 | 0.19 |
| 24.2 | 0.58 | 0.60 | 0.52 | 0.57 | 0.76 | 0.35 | 0.50 | 0.54 | 0.16 | 0.03 | 0.06 | 0.08 | 0.28 | 0.03 | 0.07 | 0.13 | 0.23 | 0.12 | 0.17 | 0.17 | 0.15 | 0.02 | 0.20 | 0.12 |
| 24.3 | 0.68 | 0.69 | 0.67 | 0.68 | 0.58 | 0.55 | 0.69 | 0.61 | 0.36 | 0.23 | 0.26 | 0.28 | 0.56 | 0.44 | 0.41 | 0.47 | 0.42 | 0.26 | 0.16 | 0.28 | 0.35 | 0.12 | 0.25 | 0.24 |
| 25.1 | 0.98 | 0.96 | 0.83 | 0.93 | 1.00 | 0.98 | 0.95 | 0.97 | 0.48 | 0.56 | 0.26 | 0.43 | 0.16 | 0.01 | 0.02 | 0.06 | 0.02 | 0.03 | 0.12 | 0.06 | 0.11 | 0.00 | 0.19 | 0.10 |
| 25.2 | 0.64 | 0.46 | 0.44 | 0.51 | 0.56 | 0.28 | 0.33 | 0.39 | 0.40 | 0.09 | 0.12 | 0.20 | 0.64 | 0.63 | 0.51 | 0.59 | 0.34 | 0.27 | 0.19 | 0.27 | 0.39 | 0.34 | 0.31 | 0.35 |
| 27 | 0.82 | 0.66 | 0.97 | 0.82 | 0.80 | 0.96 | 0.93 | 0.90 | 0.08 | 0.02 | 0.04 | 0.05 | 0.44 | 0.26 | 0.30 | 0.33 | 0.16 | 0.23 | 0.17 | 0.19 | 0.34 | 0.28 | 0.38 | 0.34 |
| 28.1 | 0.76 | 0.65 | 0.65 | 0.69 | 0.64 | 0.41 | 0.60 | 0.55 | 0.16 | 0.07 | 0.12 | 0.12 | 0.68 | 0.20 | 0.23 | 0.37 | 0.08 | 0.04 | 0.12 | 0.08 | 0.41 | 0.55 | 0.40 | 0.45 |
| 28.2 | 0.72 | 0.77 | 0.70 | 0.73 | 0.54 | 0.87 | 0.81 | 0.74 | 0.44 | 0.01 | 0.11 | 0.19 | 0.68 | 0.83 | 0.84 | 0.78 | 0.36 | 0.12 | 0.18 | 0.22 | 0.63 | 0.64 | 0.35 | 0.54 |
| 29 | 0.50 | 0.63 | 0.66 | 0.60 | 0.54 | 0.19 | 0.38 | 0.37 | 0.28 | 0.10 | 0.13 | 0.17 | 0.60 | 0.44 | 0.36 | 0.47 | 0.24 | 0.12 | 0.19 | 0.19 | 0.51 | 0.31 | 0.42 | 0.41 |
| 33 | 0.78 | 0.69 | 0.64 | 0.70 | 0.62 | 0.69 | 0.60 | 0.64 | 0.24 | 0.63 | 0.48 | 0.45 | 0.72 | 0.57 | 0.57 | 0.62 | 0.18 | 0.16 | 0.16 | 0.17 | 0.31 | 0.83 | 0.36 | 0.50 |
| 45 | 0.72 | 0.70 | 0.60 | 0.67 | 0.70 | 0.80 | 0.71 | 0.74 | 0.20 | 0.13 | 0.28 | 0.20 | 0.48 | 0.19 | 0.34 | 0.33 | 0.22 | 0.17 | 0.19 | 0.19 | 0.35 | 0.07 | 0.19 | 0.20 |

**Table S3.** List of genes in the Dipyrimicin *dip* BGC, their proposed functions, and comparison to the *cae* and *col* BGC.

| ***dip* gene** | **Size (AA)[a]** | **BLAST Hit protein [origin]** | **ID/SI[b] (%)** | ***cae* homolog** | **ID/SI[b] (%)** | ***col* homolog** | **ID/SI[b] (%)** | **proposed function** |
|---|---|---|---|---|---|---|---|---|
| *orf -3* | 123 | YciI family protein [*Actinokineospora fastidiosa*] | 90/95 | | | | | |
| *orf -2* | 425 | RNA polymerase sigma factor [Lentzea roselyniae] | 83/88 | | | | | |
| *orf -1* | 363 | FAD-dependent oxidoreductase [*Amycolatopsis japonica*] | 74/80 | | | | | |
| *1* | 201 | TetR family transcriptional regulator [*Amycolatopsis alba*] | 85/91 | *caeI2* | 33/50 | *colI2* | 30/58 | regulator |
| *2* | 135 | hypothetical protein [*Amycolatopsis sp.* WAC 04169] | 90/95 | | | | | unknown |
| *3* | 324 | glycine betaine ABC transporter substrate-binding protein [*Amycolatopsis oliviviridis*] | 96/97 | | | | | putative transporter |
| *4* | 822 | ABC transporter permease subunit [*Amycolatopsis sp.* WAC 01416] | 94/95 | | | | | putative transporter |
| *5* | 606 | asparagine synthase (glutamine hydrolyzing) [*Fodinicola feengrottensis*] | 62/73 | | | | | putative amidotransferase |
| *6* | 403 | MFS transporter [*Streptomyces sp.* NPDC004629] | 60/74 | *caeH3* | 43/60 | | | putative transporter |
| *7* | 611 | ABC transporter ATP-binding protein [*Streptomyces sp.* NPDC004629] | 68/80 | *caeH1* | 62/75 | *colH1* | 56/70 | transporter |
| *8* | 582 | ABC transporter ATP-binding protein [*Streptomyces sp.* NPDC004629] | 70/78 | *caeH2* | 63/75 | *colH2* | 59/71 | transporter |

| | | | | | | | | |
|---|---|---|---|---|---|---|---|---|
| *9* | 369 | NAD(P)/FAD-dependent oxidoreductase [*Amycolatopsis anabasis*] | 65/71 | *caeB6* | 54/66 | | | putative hydroxylase |
| *10* | 231 | alpha/beta fold hydrolase [*Streptomyces sp.* NPDC004629] | 67/76 | | | | | putative hydrolase |
| *11b* | 542 | (2,3-dihydroxybenzoyl)adenylate synthase [*Amycolatopsis anabasis*] | 73/81 | *caeA1* | 67/75 | *colA1b* | 54/63 | AMP ligase |
| *11a* | 72 | acyl carrier protein [*Micromonospora craniellae*] | 66/80 | | | *colA1a* | 46/69 | acyl carrier protein |
| *12* | 417 | DegT/DnrJ/EryC1/StrS family aminotransferase family protein [*Streptoalloteichus tenebrarius*] | 75/85 | *caeP1* | 70/80 | *colP1* | 62/74 | aminotransferase |
| *13* | 395 | FAD-dependent oxidoreductase [*Amycolatopsis anabasis*] | 76/84 | *caeP2* | 71/81 | *colP2* | 59/72 | oxidase |
| *14* | 2475 | NRPS-T1PKS [*Streptoalloteichus tenebrarius*] | 72/80 | *caeA2* | 68/77 | *colA2* | 58/69 | NRPS/PKS |
| *15* | 1058 | non-ribosomal peptide synthetase [*Amycolatopsis anabasis*] | 75/83 | *caeA3* | 71/79 | *colA3* | 52/65 | NRPS |
| *16* | 379 | acyl-CoA dehydrogenase family protein [*Amycolatopsis anabasis*] | 78/88 | *caeB1* | 79/87 | *colB1* | 62/74 | acyl-CoA dehydrogenase |
| *17* | 251 | thioesterase II family protein [*Amycolatopsis anabasis*] | 63/74 | *caeA4* | 62/72 | *colA4* | 48/63 | thioesterase |
| *18* | 145 | response regulator transcription factor [*Amycolatopsis anabasis*] | 68/76 | *caeI1* | 51/61 | *colI1* | 51/62 | regulator |
| *19* | 400 | CrmL amidohydrolase [*Actinoalloteichus sp.* WH1-2216-6] | 73/83 | *caeD* | 73/83 | *colD* | 61/73 | amidohydrolase |
| *20* | 351 | SAM-dependent methyltransferase [*Streptomyces caatingaensis*] | 65/74 | *caeG1* | 25/42 | *colG1* | 26/44 | putative O-methyltransferase |
| *21* | 550 | class I adenylate-forming enzyme family protein [*Streptomyces caatingaensis*] | 64/73 | | | | | putative AMP-dependent synthetase/ligase |
| orf 1 | 394 | MFS transporter [*Amycolatopsis sp.* NEAU-NG30] | 89/91 | | | | | |
| orf 2 | 227 | FadR/GntR family transcriptional regulator [*Streptomyces caatingaensis*] | 66/75 | | | | | |
| orf 3 | 149 | MerR family transcriptional regulator [*Amycolatopsis sp.* NEAU-NG30] | 83/88 | | | | | |

[a] denotes amino acids
[b] denotes Identity/Similarity (%)

**Table S4.** First 10 FoldSeek structural search hits for putative hydroxylase Dip9 from PDB and AlphaFold Databases.

| Target | Database | Description | Scientific Name | Prob. | Seq. Id. | TM-score | Score |
|---|---|---|---|---|---|---|---|
| 3alm | PDB100 | Crystal structure of 2-methyl-3-hydroxypyridine-5-carboxylic acid oxygenase, mutant C294A | *Mesorhizobium japonicum MAFF 303099* | 1 | 38.8 | 0.922 | 92 |
| 4h2p | PDB100 | Tetrameric form of 2-methyl-3-hydroxypyridine-5-carboxylic acid oxygenase (MHPCO) | *Mesorhizobium japonicum MAFF 303099* | 1 | 38.8 | 0.922 | 92 |
| 3all | PDB100 | Crystal structure of 2-methyl-3-hydroxypyridine-5-carboxylic acid oxygenase, mutant Y270A | *Mesorhizobium japonicum MAFF 303099* | 1 | 38.6 | 0.921 | 92 |

| | | | | | | | |
|---|---|---|---|---|---|---|---|
| 4h2n | PDB100 | Crystal structure of MHPCO, Y270F mutant | *Mesorhizobium japonicum MAFF 303099* | 1 | 38.6 | 0.921 | 92 |
| 4bk3 | PDB100 | Crystal structure of 3-hydroxybenzoate 6-hydroxylase uncovers lipid- assisted flavoprotein strategy for regioselective aromatic hydroxylation: Y105F mutant | *Rhodococcus jostii RHA1* | 0.99 | 21.9 | 0.859 | 88 |
| 4bk2 | PDB100 | Crystal structure of 3-hydroxybenzoate 6-hydroxylase uncovers lipid- assisted flavoprotein strategy for regioselective aromatic hydroxylation: Q301E mutant | *Rhodococcus jostii RHA1* | 0.99 | 22.1 | 0.858 | 88 |
| 4bjz | PDB100 | Crystal structure of 3-hydroxybenzoate 6-hydroxylase uncovers lipid- assisted flavoprotein strategy for regioselective aromatic hydroxylation: Native data | *Rhodococcus jostii RHA1* | 0.99 | 22.1 | 0.858 | 88 |
| 4bjy | PDB100 | Crystal structure of 3-hydroxybenzoate 6-hydroxylase uncovers lipid- assisted flavoprotein strategy for regioselective aromatic hydroxylation: Platinum derivative | *Rhodococcus jostii RHA1* | 0.99 | 21.9 | 0.857 | 88 |
| 3rp8 | PDB100 | Crystal Structure of Klebsiella pneumoniae R204Q HpxO complexed with FAD | *Klebsiella pneumoniae subsp. pneumoniae MGH 78578* | 0.99 | 19.4 | 0.853 | 86 |
| 4bk1 | PDB100 | Crystal structure of 3-hydroxybenzoate 6-hydroxylase uncovers lipid- assisted flavoprotein strategy for regioselective aromatic hydroxylation: H213S mutant in complex with 3-hydroxybenzoate | *Rhodococcus jostii RHA1* | 0.99 | 21.6 | 0.852 | 87 |
| AF-A0A0E4AFG7-F1-model_v4 | AFDB-SWISSPROT | Putative 2-heptyl-3-hydroxy-4(1H)-quinolone synthase AqdB1 | *Rhodococcus erythropolis* | 1 | 35.5 | 0.914 | 92 |
| AF-B1MFK1-F1-model_v4 | AFDB-SWISSPROT | 2-heptyl-3-hydroxy-4(1H)-quinolone synthase | *Mycobacteroides abscessus ATCC 19977* | 1 | 36 | 0.908 | 91 |
| AF-A0A0E4AFH6-F1-model_v4 | AFDB-SWISSPROT | Probable 2-heptyl-3-hydroxy-4(1H)-quinolone synthase AqdB2 | *Rhodococcus erythropolis* | 1 | 33.7 | 0.906 | 92 |
| AF-O06489-F1-model_v4 | AFDB-SWISSPROT | Putative oxidoreductase YetM | *Bacillus subtilis subsp. subtilis str. 168* | 0.99 | 23.7 | 0.879 | 87 |
| AF-H1ZZA4-F1-model_v4 | AFDB-SWISSPROT | Aurachin C monooxygenase/isomerase | *Stigmatella aurantiaca* | 0.99 | 22.6 | 0.869 | 88 |
| AF-Q6F6Y2-F1-model_v4 | AFDB-SWISSPROT | FAD-dependent urate hydroxylase | *Acinetobacter baylyi ADP1* | 0.99 | 19.9 | 0.863 | 88 |
| AF-Q9F131-F1-model_v4 | AFDB-SWISSPROT | 3-hydroxybenzoate 6-hydroxylase 1 | *Pseudomonas alcaligenes* | 0.99 | 22 | 0.86 | 88 |
| AF-A6T923-F1-model_v4 | AFDB-SWISSPROT | FAD-dependent urate hydroxylase | *Klebsiella pneumoniae subsp. pneumoniae MGH 78578* | 0.99 | 20 | 0.857 | 87 |

| | | | | | | | |
|---|---|---|---|---|---|---|---|
| AF-B6D1N4-F1-model_v4 | AFDB-SWISSPROT | FAD-dependent urate hydroxylase | *Klebsiella pneumoniae* | 0.99 | 20.2 | 0.856 | 87 |
| AF-B5B0J6-F1-model_v4 | AFDB-SWISSPROT | FAD-dependent urate hydroxylase | *Klebsiella oxytoca* | 0.99 | 17.9 | 0.855 | 87 |
| AF-A0A353BC83-F1-model_v4 | AFDB50 | FAD-dependent oxidoreductase | *Planctomycetaceae bacterium* | 1 | 24.1 | 0.876 | 89 |
| AF-A0A1K1MJG4-F1-model_v4 | AFDB50 | 2-polyprenyl-6-methoxyphenol hydroxylase | *Luteibacter sp. UNCMF366Tsu5.1* | 0.99 | 25.1 | 0.875 | 88 |
| AF-A0A521KTI3-F1-model_v4 | AFDB50 | FAD-dependent monooxygenase | *Nitrospirae bacterium* | 0.99 | 23 | 0.875 | 88 |
| AF-A0A5C6BNZ2-F1-model_v4 | AFDB50 | FAD-dependent urate hydroxylase | *Symmachiella macrocystis* | 0.99 | 21.3 | 0.873 | 88 |
| AF-A0A0A3HX39-F1-model_v4 | AFDB50 | FAD_binding_3 domain-containing protein | *Ureibacillus sinduriensis BLB-1 = JCM 15800* | 0.99 | 19 | 0.873 | 88 |
| AF-A0A168HQA1-F1-model_v4 | AFDB50 | FAD_binding_3 domain-containing protein | *Paenibacillus glacialis* | 1 | 19.5 | 0.873 | 89 |
| AF-A0A7W1I6S8-F1-model_v4 | AFDB50 | FAD-dependent monooxygenase | *Actinomycetia bacterium* | 0.99 | 22.3 | 0.872 | 88 |
| AF-A0A4R6ULI2-F1-model_v4 | AFDB50 | 2-polyprenyl-6-methoxyphenol hydroxylase-like FAD-dependent oxidoreductase | *Permianibacter aggregans* | 0.99 | 18.6 | 0.872 | 86 |
| AF-A0A379BP18-F1-model_v4 | AFDB50 | 3-hydroxybenzoate 6-hydroxylase 1 | *Nocardia brasiliensis* | 0.99 | 21.6 | 0.872 | 87 |
| AF-A0A1H7VYS7-F1-model_v4 | AFDB50 | 2-polyprenyl-6-methoxyphenol hydroxylase | *Stigmatella aurantiaca* | 0.99 | 22.3 | 0.871 | 88 |
| AF-K0EM28-F1-model_v4 | AFDB-PROTEOME | Monooxygenase | *Nocardia brasiliensis ATCC 700358* | 0.99 | 25.4 | 0.872 | 86 |
| AF-K0ETR9-F1-model_v4 | AFDB-PROTEOME | FAD-binding monooxygenase | *Nocardia brasiliensis ATCC 700358* | 0.99 | 21.8 | 0.871 | 87 |
| AF-Q9HYR7-F1-model_v4 | AFDB-PROTEOME | Probable FAD-dependent monooxygenase | *Pseudomonas aeruginosa PAO1* | 0.99 | 19.3 | 0.866 | 88 |

| | | | | | | | |
|---|---|---|---|---|---|---|---|
| AF-Q2FVW3-F1-model_v4 | AFDB-PROTEOME | FAD_binding_3 domain-containing protein | *Staphylococcus aureus subsp. aureus NCTC 8325* | 0.99 | 21 | 0.864 | 86 |
| AF-K0ETS7-F1-model_v4 | AFDB-PROTEOME | 2-polyprenyl-6-methoxyphenol hydroxylase-like oxidoreductase | *Nocardia brasiliensis ATCC 700358* | 0.99 | 21.3 | 0.861 | 86 |
| AF-A0A0H3GN89-F1-model_v4 | AFDB-PROTEOME | Putative flavoprotein monooxygenase | *Klebsiella pneumoniae subsp. pneumoniae HS11286* | 0.99 | 20 | 0.857 | 87 |
| AF-K0ELV8-F1-model_v4 | AFDB-PROTEOME | FAD-binding monooxygenase | *Nocardia brasiliensis ATCC 700358* | 0.99 | 21.5 | 0.847 | 86 |
| AF-Q10RL6-F1-model_v4 | AFDB-PROTEOME | FAD binding domain containing protein, expressed | *Oryza sativa Japonica Group* | 0.99 | 19 | 0.847 | 88 |
| AF-Q9FLC2-F1-model_v4 | AFDB-PROTEOME | Monooxygenase 3 | *Arabidopsis thaliana* | 0.99 | 18.4 | 0.847 | 88 |
| AF-O81816-F1-model_v4 | AFDB-PROTEOME | Monooxygenase 2 | *Arabidopsis thaliana* | 0.99 | 19.6 | 0.847 | 88 |

**Table S5.** First 10 FoldSeek structural search hits for putative O-methyltransferae Dip20 from PDB and AlphaFold Databases.

| Target | Database | Description | Scientific Name | Prob. | Seq. Id. | TM-score | Score |
|---|---|---|---|---|---|---|---|
| 2r3s | PDB100 | Crystal structure of a putative O-methyltransferase at 2.15 A resolution | *Nostoc punctiforme PCC 73102* | 0.99 | 26.1 | 0.907 | 87 |
| 7clf | PDB100 | PigF with SAH | *Serratia marcescens* | 0.99 | 21 | 0.853 | 83 |
| 4a6d | PDB100 | Crystal structure of human N-acetylserotonin methyltransferase (ASMT) in complex with SAM | *Homo sapiens* | 0.99 | 22.1 | 0.848 | 84 |
| 6c5b | PDB100 | Crystal Structure Analysis of LaPhzM | *Lysobacter antibioticus* | 0.99 | 18.7 | 0.84 | 81 |
| 7wdq | PDB100 | DsyB in complex with SAM | *Nisaea denitrificans DSM 18348* | 0.99 | 22.7 | 0.835 | 81 |
| 8tjj | PDB100 | SAM-dependent methyltransferase RedM bound to SAM | *uncultured bacterium* | 0.98 | 24.6 | 0.831 | 79 |
| 5icc | PDB100 | Crystal structure of (S)-norcoclaurine 6-O-methyltransferase with S-adenosyl-L-homocysteine | *Thalictrum flavum subsp. glaucum* | 0.99 | 16.8 | 0.829 | 83 |
| 7v6l | PDB100 | LcCOMT in complex with SAH | Ligusticum chuanxiong | 0.99 | 16.8 | 0.827 | 81 |
| 3gwz | PDB100 | Structure of the Mitomycin 7-O-methyltransferase MmcR | Streptomyces lavendulae | 0.99 | 22.7 | 0.827 | 81 |
| 6yjw | PDB100 | Structure of Fragaria ananassa O-methyltransferase crystallized with PAS polypeptide | Fragaria x ananassa | 0.99 | 17.7 | 0.824 | 82 |
| AF-A0A7Y5HBN6-F1-model_v4 | AFDB50 | Methyltransferase domain-containing protein | *Candidatus Brocadiae bacterium* | 1 | 34.6 | 0.922 | 90 |

| | | | | | | | |
|---|---|---|---|---|---|---|---|
| AF-A0A1E4G6R2-F1-model_v4 | AFDB50 | Uncharacterized protein | *bacterium SCN 62-11* | 0.99 | 26.5 | 0.872 | 84 |
| AF-A0A2S6QMZ9-F1-model_v4 | AFDB50 | 3-hydroxy-5-methyl-1-naphthoate 3-O-methyltransferase | *Alphaproteobacteria bacterium MarineAlpha9_Bin3* | 0.99 | 18.9 | 0.866 | 85 |
| AF-A0A5J6GJ07-F1-model_v4 | AFDB50 | Methyltransf_2 domain-containing protein | *Streptomyces kanamyceticus* | 0.99 | 22.2 | 0.863 | 83 |
| AF-A0A2R5HC89-F1-model_v4 | AFDB50 | O-methyltransferase | *Mycobacterium montefiorense* | 0.99 | 20.9 | 0.857 | 83 |
| AF-A0A254QRY0-F1-model_v4 | AFDB50 | Methyltransferase | *Phaeobacter sp. 22II1-1F12B* | 0.99 | 20.9 | 0.854 | 83 |
| AF-A0A7C2P238-F1-model_v4 | AFDB50 | Methyltransferase domain-containing protein | *Schlesneria paludicola* | 0.99 | 20.9 | 0.852 | 84 |
| AF-A0A1H1GZC4-F1-model_v4 | AFDB50 | Dimerisation domain-containing protein | *Actinopolyspora saharensis* | 0.99 | 20.6 | 0.851 | 84 |
| AF-A0A501PB90-F1-model_v4 | AFDB50 | Methyltransferase | *Emcibacter nanhaiensis* | 0.99 | 14.2 | 0.849 | 82 |
| AF-A0A7W8JHX0-F1-model_v4 | AFDB50 | Putative transcriptional regulator | *Rhodanobacter sp. ANJX3* | 0.99 | 17.9 | 0.849 | 83 |
| AF-Q92056-F1-model_v4 | AFDB-SWISSPROT | Acetylserotonin O-methyltransferase | *Gallus gallus* | 0.99 | 20.9 | 0.86 | 85 |
| AF-P10950-F1-model_v4 | AFDB-SWISSPROT | Acetylserotonin O-methyltransferase | *Bos taurus* | 0.99 | 21.4 | 0.853 | 84 |
| AF-Q8HZJ0-F1-model_v4 | AFDB-SWISSPROT | Acetylserotonin O-methyltransferase | *Macaca mulatta* | 0.99 | 20.9 | 0.853 | 84 |
| AF-P46597-F1-model_v4 | AFDB-SWISSPROT | Acetylserotonin O-methyltransferase | *Homo sapiens* | 0.99 | 21.7 | 0.848 | 84 |
| AF-A0A0N9HTA1-F1-model_v4 | AFDB-SWISSPROT | Desmethyl-yatein O-methyltransferase | *Sinopodophyllum hexandrum* | 0.99 | 18.1 | 0.844 | 84 |
| AF-P39896-F1-model_v4 | AFDB-SWISSPROT | Tetracenomycin polyketide synthesis 8-O-methyl transferase TcmO | *Streptomyces glaucescens* | 0.99 | 23.4 | 0.844 | 83 |
| AF-Q42654-F1-model_v4 | AFDB-SWISSPROT | Flavonoid 3'-O-methyltransferase FOMT | *Chrysosplenium americanum* | 0.99 | 15.8 | 0.838 | 82 |
| AF-Q54B60-F1-model_v4 | AFDB-SWISSPROT | Probable inactive O-methyltransferase 11 | *Dictyostelium discoideum* | 0.99 | 14 | 0.838 | 81 |

| | | | | | | | |
|---|---|---|---|---|---|---|---|
| AF-Q42653-F1-model_v4 | AFDB-SWISSPROT | Flavone 3'-O-methyltransferase OMT2 | *Chrysosplenium americanum* | 0.99 | 16.9 | 0.83 | 82 |
| AF-Q6WUC1-F1-model_v4 | AFDB-SWISSPROT | (RS)-norcoclaurine 6-O-methyltransferase | *Papaver somniferum* | 0.99 | 14.6 | 0.828 | 82 |
| AF-P46597-F1-model_v4 | AFDB-PROTEOME | Acetylserotonin O-methyltransferase | *Homo sapiens* | 0.99 | 21.7 | 0.848 | 84 |
| AF-K0F7F0-F1-model_v4 | AFDB-PROTEOME | O-methyltransferase | *Nocardia brasiliensis ATCC 700358* | 0.99 | 21.3 | 0.847 | 85 |
| AF-A0A0R4ITX8-F1-model_v4 | AFDB-PROTEOME | Acetylserotonin O-methyltransferase | *Danio rerio* | 0.99 | 20.9 | 0.846 | 83 |
| AF-K0ESM0-F1-model_v4 | AFDB-PROTEOME | O-demethylpuromycin-O-methyltransferase | *Nocardia brasiliensis ATCC 700358* | 0.99 | 21.2 | 0.839 | 84 |
| AF-A3KNM1-F1-model_v4 | AFDB-PROTEOME | Acetylserotonin O-methyltransferase 2 | *Danio rerio* | 0.99 | 20.8 | 0.838 | 83 |
| AF-Q54B60-F1-model_v4 | AFDB-PROTEOME | Probable inactive O-methyltransferase 11 | *Dictyostelium discoideum* | 0.99 | 14 | 0.838 | 81 |
| AF-K0EN01-F1-model_v4 | AFDB-PROTEOME | O-methyltransferase family protein | *Nocardia brasiliensis ATCC 700358* | 0.99 | 26.2 | 0.834 | 82 |
| AF-Q86I40-F1-model_v4 | AFDB-PROTEOME | O-methyltransferase 4 | *Dictyostelium discoideum* | 0.99 | 12.8 | 0.83 | 81 |
| AF-O53764-F1-model_v4 | AFDB-PROTEOME | Probable methyltransferase/methylase | *Mycobacterium tuberculosis H37Rv* | 0.99 | 21.9 | 0.829 | 81 |
| AF-A0A0R4J3I5-F1-model_v4 | AFDB-PROTEOME | Uncharacterized protein | *Glycine max* | 0.99 | 17.7 | 0.822 | 83 |

**Table S6.** First 10 FoldSeek structural search hits for putative amidotransferase Dip5 from PDB and AlphaFold Databases.

| Target | Database | Description | Scientific Name | Prob. | Seq. Id. | TM-score | Score |
|---|---|---|---|---|---|---|---|
| 7ylz | PDB100 | Unliganded form of hydroxyamidotransferase TsnB9 | *Streptomyces sp. RM72* | 1 | 37.5 | 0.937 | 91 |
| 1ct9 | PDB100 | crystal structure of asparagine synthetase B from Escherichia coli | *Escherichia coli* | 0.96 | 20.5 | 0.818 | 73 |
| 6gq3 | PDB100 | Human asparagine synthetase (ASNS) in complex with 6-diazo-5-oxo-L-norleucine (DON) at 1.85 A resolution | *Homo sapiens* | 0.96 | 20.5 | 0.798 | 73 |
| 1mc1 | PDB100 | beta-lactam synthetase with product (DGPC), AMP and PPI | *Streptomyces clavuligerus* | 0.92 | 16.3 | 0.759 | 69 |
| 1mbz | PDB100 | beta-lactam synthetase with trapped intermediate | *Streptomyces clavuligerus* | 0.91 | 16.3 | 0.758 | 68 |
| 1mb9 | PDB100 | beta-lactam synthetase complexed with ATP | *Streptomyces clavuligerus* | 0.91 | 16.3 | 0.757 | 68 |

| | | | | | | | |
|---|---|---|---|---|---|---|---|
| 1jgt | PDB100 | crystal structure of beta-lactam synthetase | *Streptomyces clavuligerus* | 0.91 | 16.4 | 0.754 | 68 |
| 1m1z | PDB100 | beta-lactam synthetase apo enzyme | *Streptomyces clavuligerus* | 0.9 | 16.4 | 0.753 | 67 |
| 1q19 | PDB100 | Carbapenam Synthetase | *Pectobacterium carotovorum* | 0.91 | 11.7 | 0.746 | 68 |
| 1q15 | PDB100 | Carbapenam Synthetase | *Pectobacterium carotovorum* | 0.89 | 11.5 | 0.738 | 66 |
| AF-Q9I231-F1-model_v4 | AFDB-PROTEOME | Probable asparagine synthetase | *Pseudomonas aeruginosa PAO1* | 1 | 45 | 0.971 | 97 |
| AF-K0EYT1-F1-model_v4 | AFDB-PROTEOME | Putative amidotransferase | *Nocardia brasiliensis ATCC 700358* | 1 | 45.8 | 0.964 | 97 |
| AF-Q9I781-F1-model_v4 | AFDB-PROTEOME | Potential phenazine-modifying enzyme | *Pseudomonas aeruginosa PAO1* | 1 | 40.7 | 0.96 | 96 |
| AF-K0FD31-F1-model_v4 | AFDB-PROTEOME | Asparagine synthetase | *Nocardia brasiliensis ATCC 700358* | 1 | 43.3 | 0.956 | 96 |
| AF-Q86A01-F1-model_v4 | AFDB-PROTEOME | Asparagine synthetase | *Dictyostelium discoideum* | 1 | 32.8 | 0.934 | 96 |
| AF-Q9HYE7-F1-model_v4 | AFDB-PROTEOME | Probable glutamine amidotransferase | *Pseudomonas aeruginosa PAO1* | 1 | 29 | 0.902 | 89 |
| AF-A0A132Z0T1-F1-model_v4 | AFDB-PROTEOME | Asparagine synthase (Glutamine-hydrolyzing) | *Enterococcus faecium* | 0.99 | 24 | 0.867 | 88 |
| AF-P9WN33-F1-model_v4 | AFDB-PROTEOME | Putative asparagine synthetase [glutamine-hydrolyzing] | *Mycobacterium tuberculosis H37Rv* | 0.99 | 24.1 | 0.858 | 88 |
| AF-K0EXR4-F1-model_v4 | AFDB-PROTEOME | Asparagine synthase | *Nocardia brasiliensis ATCC 700358* | 0.99 | 22.1 | 0.85 | 88 |
| AF-Q9CCF2-F1-model_v4 | AFDB-PROTEOME | Putative asparagine synthetase | *Mycobacterium leprae TN* | 0.99 | 23.2 | 0.845 | 88 |
| AF-A0A0K0QAS6-F1-model_v4 | AFDB50 | Asparagine synthase | *Bacillus thuringiensis* | 1 | 41.8 | 0.971 | 97 |
| AF-A0A0B5NI93-F1-model_v4 | AFDB50 | Asparagine synthase (Glutamine-hydrolyzing) | *Bacillus thuringiensis* | 1 | 42.2 | 0.971 | 97 |
| AF-A0A854KS77-F1-model_v4 | AFDB50 | Asparagine synthase (Glutamine-hydrolyzing) | *Bacillus thuringiensis serovar shandongiensis* | 1 | 42.4 | 0.971 | 97 |
| AF-A0A090YNR9-F1-model_v4 | AFDB50 | Asparagine synthase | *Bacillus clarus* | 1 | 42.1 | 0.971 | 97 |
| AF-A0A373KWD5-F1-model_v4 | AFDB50 | Asparagine synthase (Glutamine-hydrolyzing) | *Roseburia sp. AF42-8* | 1 | 36.2 | 0.969 | 97 |
| AF-R5JEM6-F1-model_v4 | AFDB50 | Asparagine synthetase | *Coprococcus sp. CAG:782* | 1 | 33.3 | 0.968 | 97 |

| | | | | | | | |
|---|---|---|---|---|---|---|---|
| AF-R7RSB4-F1-model_v4 | AFDB50 | Asparagine synthetase [glutamine-hydrolyzing] | *Thermobrachium celere DSM 8682* | 1 | 34.9 | 0.968 | 97 |
| AF-A0A6G2UGB8-F1-model_v4 | AFDB50 | Asparagine synthase (Glutamine-hydrolyzing) | *Streptomyces sp. SID4931* | 1 | 46.4 | 0.967 | 97 |
| AF-A0A3M8H280-F1-model_v4 | AFDB50 | Asparagine synthase (Glutamine-hydrolyzing) | *Lysinibacillus halotolerans* | 1 | 39.6 | 0.967 | 97 |
| AF-A0A7K0CDT8-F1-model_v4 | AFDB50 | Asparagine synthetase [glutamine-hydrolyzing] 3 | *Streptomyces smaragdinus* | 1 | 45.5 | 0.962 | 97 |
| AF-O05272-F1-model_v4 | AFDB-SWISSPROT | Asparagine synthetase [glutamine-hydrolyzing] 3 | *Bacillus subtilis subsp. subtilis str. 168* | 1 | 40.7 | 0.968 | 97 |
| AF-P54420-F1-model_v4 | AFDB-SWISSPROT | Asparagine synthetase [glutamine-hydrolyzing] 1 | *Bacillus subtilis subsp. subtilis str. 168* | 0.99 | 25.8 | 0.872 | 88 |
| AF-P9WN33-F1-model_v4 | AFDB-SWISSPROT | Putative asparagine synthetase [glutamine-hydrolyzing] | *Mycobacterium tuberculosis H37Rv* | 0.99 | 24.1 | 0.858 | 88 |
| AF-P64248-F1-model_v4 | AFDB-SWISSPROT | Putative asparagine synthetase [glutamine-hydrolyzing] | *Mycobacterium tuberculosis variant bovis AF2122/97* | 0.99 | 24.1 | 0.858 | 88 |
| AF-P9WN32-F1-model_v4 | AFDB-SWISSPROT | Putative asparagine synthetase [glutamine-hydrolyzing] | *Mycobacterium tuberculosis CDC1551* | 0.99 | 24 | 0.858 | 88 |
| AF-B6HLP8-F1-model_v4 | AFDB-SWISSPROT | Amidase chyE | *Penicillium rubens Wisconsin 54-1255* | 1 | 19.4 | 0.855 | 89 |
| AF-I1S3K8-F1-model_v4 | AFDB-SWISSPROT | Amidase chry2 | *Fusarium graminearum PH-1* | 1 | 19.6 | 0.855 | 89 |
| AF-O24338-F1-model_v4 | AFDB-SWISSPROT | Asparagine synthetase [glutamine-hydrolyzing] | *Sandersonia aurantiaca* | 0.96 | 21.5 | 0.804 | 74 |
| AF-Q54MB4-F1-model_v4 | AFDB-SWISSPROT | Probable asparagine synthetase [glutamine-hydrolyzing] | *Dictyostelium discoideum* | 0.97 | 20.8 | 0.785 | 75 |
| AF-P22106-F1-model_v4 | AFDB-SWISSPROT | Asparagine synthetase B [glutamine-hydrolyzing] | *Escherichia coli K-12* | 0.96 | 21.1 | 0.781 | 74 |

**Table S7.** First 10 FoldSeek structural search hits for putative synthetase/ligase Dip21 from PDB and AlphaFold Databases.

| **Target** | **Database** | **Description** | **Scientific Name** | **Prob.** | **Seq. Id.** | **TM-score** | **Score** |
|---|---|---|---|---|---|---|---|
| 1mdf | PDB100 | Crystal structure of DhbE in absence of substrate | *Bacillus subtilis* | 1 | 28.7 | 0.913 | 90 |
| 7tz4 | PDB100 | Salicylate Adenylate PchD from Pseudomonas aeruginosa containing 4-cyanosalicyl-AMS | *Pseudomonas aeruginosa* | 0.99 | 28.8 | 0.899 | 88 |
| 7tyb | PDB100 | Salicylate Adenylate PchD from Pseudomonas aeruginosa containing salicyl-AMS | *Pseudomonas aeruginosa PAO1* | 0.99 | 28.8 | 0.895 | 87 |
| 7kyd | PDB100 | Drosophila melanogaster long-chain fatty-acyl-CoA synthetase | *Drosophila melanogaster* | 0.99 | 20.4 | 0.869 | 85 |

| | | | | | | | |
|---|---|---|---|---|---|---|---|
| | | CG6178 | | | | | |
| 4gxr | PDB100 | Structure of ATP bound RpMatB-BxBclM chimera B3 | *Rhodopseudomonas palustris CGA009* | 0.99 | 22.5 | 0.869 | 83 |
| 2v7b | PDB100 | Crystal structures of a benzoate CoA ligase from Burkholderia xenovorans LB400 | *Paraburkholderia xenovorans LB400* | 0.99 | 20.2 | 0.867 | 81 |
| 4gxq | PDB100 | Crystal Structure of ATP bound RpMatB-BxBclM chimera B1 | *Rhodopseudomonas palustris CGA009* | 0.99 | 23.2 | 0.866 | 83 |
| 4wv3 | PDB100 | Crystal structure of the anthranilate CoA ligase AuaEII in complex with anthranoyl-AMP | *Stigmatella aurantiaca* | 0.99 | 21.3 | 0.866 | 84 |
| 5bsw | PDB100 | Crystal structure of 4-coumarate:CoA ligase delta-V341 mutant complexed with feruloyl adenylate | *Nicotiana tabacum* | 0.99 | 21.3 | 0.862 | 84 |
| 6h1b | PDB100 | Structure of amide bond synthetase Mcba K483A mutant from Marinactinospora thermotolerans | *Marinactinospora thermotolerans* | 0.99 | 28.8 | 0.861 | 81 |
| AF-A0A534XEG6-F1-model_v4 | AFDB50 | Cyclohexanecarboxylate-CoA ligase | *Deltaproteobacteria bacterium* | 1 | 30.4 | 0.924 | 92 |
| AF-A0A7Y0JBY0-F1-model_v4 | AFDB50 | Acyl--CoA ligase | *Streptomyces sp. GMY02* | 1 | 39.2 | 0.918 | 90 |
| AF-A0A6J7NLY6-F1-model_v4 | AFDB50 | Unannotated protein | *freshwater metagenome* | 1 | 26.3 | 0.905 | 90 |
| AF-A0A1C5LZ60-F1-model_v4 | AFDB50 | Short-chain-fatty-acid--CoA ligase | *uncultured Clostridium sp.* | 1 | 24.8 | 0.902 | 90 |
| AF-A0A1I7GUI0-F1-model_v4 | AFDB50 | Cyclohexanecarboxylate-CoA ligase | *Alicyclobacillus macrosporangiidus* | 1 | 28.2 | 0.896 | 89 |
| AF-A0A7C2Q990-F1-model_v4 | AFDB50 | Long-chain fatty acid--CoA ligase | *Armatimonadetes bacterium* | 0.99 | 29.2 | 0.893 | 86 |
| AF-A0A534X5V8-F1-model_v4 | AFDB50 | Cyclohexanecarboxylate-CoA ligase | *Deltaproteobacteria bacterium* | 0.99 | 31 | 0.892 | 86 |
| AF-A0A6J7NHN7-F1-model_v4 | AFDB50 | Unannotated protein | *freshwater metagenome* | 0.99 | 23.3 | 0.888 | 88 |
| AF-A0A7G3AAM8-F1-model_v4 | AFDB50 | Putative acyl-coa synthetase family member 2 isoform x2 | *Lutzomyia longipalpis* | 0.99 | 23.3 | 0.886 | 88 |
| AF-A0A235HXW3-F1-model_v4 | AFDB50 | Long-chain fatty acid--CoA ligase | *Nostoc sp. 'Peltigera membranacea cyanobiont' 210A* | 0.99 | 25.6 | 0.883 | 84 |

| | | | | | | | |
|---|---|---|---|---|---|---|---|
| AF-K0EMM1-F1-model_v4 | AFDB-PROTEOME | Hydroxybenzoate-AMP ligase | *Nocardia brasiliensis ATCC 700358* | 1 | 28.7 | 0.919 | 90 |
| AF-K0ER74-F1-model_v4 | AFDB-PROTEOME | (2,3-dihydroxybenzoyl)adenylate synthase | *Nocardia brasiliensis ATCC 700358* | 1 | 27.3 | 0.895 | 89 |
| AF-K0F5E8-F1-model_v4 | AFDB-PROTEOME | Uncharacterized protein | *Nocardia brasiliensis ATCC 700358* | 0.99 | 28 | 0.882 | 85 |
| AF-I6Y0X0-F1-model_v4 | AFDB-PROTEOME | Probable fatty-acid-CoA ligase FadD35 (Fatty-acid-CoA synthetase) (Fatty-acid-CoA synthase) | *Mycobacterium tuberculosis H37Rv* | 0.99 | 28.2 | 0.88 | 87 |
| AF-P31552-F1-model_v4 | AFDB-PROTEOME | Crotonobetaine/carnitine--CoA ligase | *Escherichia coli K-12* | 0.99 | 19.1 | 0.874 | 84 |
| AF-Q9I0S7-F1-model_v4 | AFDB-PROTEOME | Probable AMP-binding enzyme | *Pseudomonas aeruginosa PAO1* | 0.99 | 26 | 0.872 | 88 |
| AF-Q8ZRX4-F1-model_v4 | AFDB-PROTEOME | Crotonobetaine/carnitine--CoA ligase | *Salmonella enterica subsp. enterica serovar Typhimurium str. LT2* | 0.99 | 19.3 | 0.872 | 84 |
| AF-C1GZT6-F1-model_v4 | AFDB-PROTEOME | Short-chain-fatty-acid-CoA ligase | *Paracoccidioides lutzii Pb01* | 0.99 | 22 | 0.871 | 88 |
| AF-Q9VDU4-F1-model_v4 | AFDB-PROTEOME | Uncharacterized protein | *Drosophila melanogaster* | 0.99 | 15.7 | 0.87 | 86 |
| AF-Q5F969-F1-model_v4 | AFDB-PROTEOME | Long-chain fatty acid--CoA ligase | *Neisseria gonorrhoeae FA 1090* | 0.99 | 21.9 | 0.867 | 84 |
| AF-Q84HC5-F1-model_v4 | AFDB-SWISSPROT | 2-hydroxy-7-methoxy-5-methyl-1-naphthoate--CoA ligase | *Streptomyces carzinostaticus* | 1 | 28.7 | 0.91 | 91 |
| AF-P40871-F1-model_v4 | AFDB-SWISSPROT | 2,3-dihydroxybenzoate-AMP ligase | *Bacillus subtilis subsp. subtilis str. 168* | 1 | 28.2 | 0.909 | 89 |
| AF-Q7X279-F1-model_v4 | AFDB-SWISSPROT | Salicylyl-CoA synthase / salicylate adenylyltransferase | *Streptomyces sp.* | 1 | 26.5 | 0.898 | 90 |
| AF-Q0TLV2-F1-model_v4 | AFDB-SWISSPROT | Crotonobetaine/carnitine--CoA ligase | *Escherichia coli 536* | 0.99 | 20.2 | 0.888 | 86 |
| AF-B4TWR4-F1-model_v4 | AFDB-SWISSPROT | Crotonobetaine/carnitine--CoA ligase | *Salmonella enterica subsp. enterica serovar Schwarzengrund str. CVM19633* | 0.99 | 20.8 | 0.885 | 85 |
| AF-B5F750-F1-model_v4 | AFDB-SWISSPROT | Crotonobetaine/carnitine--CoA ligase | *Salmonella enterica subsp. enterica serovar Agona str. SL483* | 0.99 | 20.6 | 0.885 | 85 |
| AF-C0Q4L3-F1-model_v4 | AFDB-SWISSPROT | Crotonobetaine/carnitine--CoA ligase | *Salmonella enterica subsp. enterica serovar Paratyphi C str. RKS4594* | 0.99 | 20.4 | 0.884 | 85 |
| AF-B5R1R0-F1-model_v4 | AFDB-SWISSPROT | Crotonobetaine/carnitine--CoA ligase | *Salmonella enterica subsp. enterica serovar Enteritidis str. P125109* | 0.99 | 20.6 | 0.884 | 85 |
| AF-O07610-F1-model_v4 | AFDB-SWISSPROT | Long-chain-fatty-acid--CoA ligase | *Bacillus subtilis subsp. subtilis str. 168* | 0.99 | 23.3 | 0.882 | 85 |
| AF-O31826-F1-model_v4 | AFDB-SWISSPROT | Putative acyl-CoA synthetase YngI | *Bacillus subtilis subsp. subtilis str. 168* | 0.99 | 23.5 | 0.882 | 88 |

**Table S8.** First 10 FoldSeek structural search hits for putative hydrolase Dip10 from PDB and AlphaFold Databases.

| Target | Database | Description | Scientific Name | Prob. | Seq. Id. | TM-score | Score |
|---|---|---|---|---|---|---|---|
| 4l0c | PDB100 | Crystal structure of the N-Fopmylmaleamic acid deformylase Nfo(S94A) from Pseudomonas putida S16 | *Pseudomonas putida S16* | 0.99 | 19.7 | 0.817 | 84 |
| 3kxp | PDB100 | Crystal Structure of E-2-(Acetamidomethylene)succinate Hydrolase | *Mesorhizobium loti* | 0.99 | 19.6 | 0.805 | 85 |
| 1zoi | PDB100 | Crystal Structure of a Stereoselective Esterase from Pseudomonas putida IFO12996 | *Pseudomonas putida* | 0.99 | 15.7 | 0.794 | 85 |
| 1a88 | PDB100 | CHLOROPEROXIDASE L | *Streptomyces lividans* | 0.99 | 15.3 | 0.793 | 85 |
| 5h3h | PDB100 | Esterase (EaEST) from Exiguobacterium antarcticum | *Exiguobacterium antarcticum B7* | 0.99 | 14.2 | 0.791 | 84 |
| 3hea | PDB100 | The L29P/L124I mutation of Pseudomonas fluorescens esterase | *Pseudomonas fluorescens* | 0.99 | 13.6 | 0.791 | 84 |
| 3hi4 | PDB100 | Switching catalysis from hydrolysis to perhydrolysis in P. fluorescens esterase | *Pseudomonas fluorescens* | 0.99 | 13.6 | 0.79 | 84 |
| 3ia2 | PDB100 | Pseudomonas fluorescens esterase complexed to the R-enantiomer of a sulfonate transition state analog | *Pseudomonas fluorescens* | 0.99 | 13.3 | 0.79 | 84 |
| 3t4u | PDB100 | L29I Mutation in an Aryl Esterase from Pseudomonas fluorescens Leads to Unique Peptide Flip and Increased Activity | *Pseudomonas fluorescens* | 0.99 | 13.6 | 0.79 | 84 |
| 4dgq | PDB100 | Crystal structure of Non-heme chloroperoxidase from Burkholderia cenocepacia | *Burkholderia cenocepacia J2315* | 0.99 | 14.2 | 0.787 | 85 |
| AF-M2QRQ5-F1-model_v4 | AFDB50 | Putative hydrolase | *Amycolatopsis azurea DSM 43854* | 1 | 99.5 | 0.949 | 91 |
| AF-A0A7K0DV75-F1-model_v4 | AFDB50 | 2-succinyl-6-hydroxy-2, 4-cyclohexadiene-1-carboxylate synthase | *Nocardia aurantia* | 1 | 46.3 | 0.947 | 94 |
| AF-A0A7H1Q9C4-F1-model_v4 | AFDB50 | Alpha/beta hydrolase | *Streptomyces griseofuscus* | 1 | 40 | 0.946 | 94 |
| AF-A0A426R654-F1-model_v4 | AFDB50 | Alpha/beta fold hydrolase | *Rhodococcus sp. Eu-32* | 1 | 46.9 | 0.946 | 94 |
| AF-A0A3A4AC57-F1-model_v4 | AFDB50 | Alpha/beta fold hydrolase | *Bailinhaonella thermotolerans* | 1 | 43.1 | 0.938 | 94 |
| AF-A0A6M4PRH7-F1-model_v4 | AFDB50 | Alpha/beta hydrolase | *Streptomyces sp. Jing01* | 1 | 42.6 | 0.935 | 93 |
| AF-A0A1G8TD96-F1- | AFDB50 | Alpha/beta hydrolase family protein | *Streptomyces indicus* | 1 | 37.7 | 0.934 | 93 |

| | | | | | | | |
|---|---|---|---|---|---|---|---|
| model_v4 | | | | | | | |
| AF-A0A387BNR8-F1-model_v4 | AFDB50 | Alpha/beta hydrolase | *Gryllotalpicola protaetiae* | 1 | 41.3 | 0.93 | 94 |
| AF-A0A652KQZ7-F1-model_v4 | AFDB50 | Alpha/beta fold hydrolase | *Streptomyces sp. adm13(2018)* | 1 | 40.2 | 0.93 | 93 |
| AF-A0A6L9Y0K9-F1-model_v4 | AFDB50 | Alpha/beta hydrolase | *Diaminobutyricibacter tongyongensis* | 1 | 41.4 | 0.929 | 93 |
| AF-Q88FY3-F1-model_v4 | AFDB-SWISSPROT | N-formylmaleamate deformylase | *Pseudomonas putida KT2440* | 0.99 | 17 | 0.812 | 86 |
| AF-Q609V0-F1-model_v4 | AFDB-SWISSPROT | Pimeloyl-[acyl-carrier protein] methyl ester esterase | *Methylococcus capsulatus str. Bath* | 0.99 | 16 | 0.805 | 83 |
| AF-B5FFE9-F1-model_v4 | AFDB-SWISSPROT | Pimeloyl-[acyl-carrier protein] methyl ester esterase | *Aliivibrio fischeri MJ11* | 0.99 | 13.8 | 0.8 | 83 |
| AF-Q5E8N3-F1-model_v4 | AFDB-SWISSPROT | Pimeloyl-[acyl-carrier protein] methyl ester esterase | *Aliivibrio fischeri ES114* | 0.99 | 13.5 | 0.799 | 83 |
| AF-A7MST3-F1-model_v4 | AFDB-SWISSPROT | Pimeloyl-[acyl-carrier protein] methyl ester esterase | *Vibrio campbellii CAIM 519 = NBRC 15631 = ATCC 25920* | 0.99 | 15.6 | 0.798 | 83 |
| AF-A0KF11-F1-model_v4 | AFDB-SWISSPROT | Pimeloyl-[acyl-carrier protein] methyl ester esterase | *Aeromonas hydrophila subsp. hydrophila ATCC 7966* | 0.99 | 19.3 | 0.797 | 83 |
| AF-O32234-F1-model_v4 | AFDB-SWISSPROT | AB hydrolase superfamily protein YvaM | *Bacillus subtilis subsp. subtilis str. 168* | 0.99 | 13.2 | 0.797 | 83 |
| AF-A4ST17-F1-model_v4 | AFDB-SWISSPROT | Pimeloyl-[acyl-carrier protein] methyl ester esterase | *Aeromonas salmonicida subsp. salmonicida A449* | 0.99 | 17.3 | 0.796 | 82 |
| AF-O05691-F1-model_v4 | AFDB-SWISSPROT | Non-heme haloperoxidase | *Rhodococcus erythropolis* | 0.99 | 16.5 | 0.796 | 85 |
| AF-Q5QZC0-F1-model_v4 | AFDB-SWISSPROT | Pimeloyl-[acyl-carrier protein] methyl ester esterase | *Idiomarina loihiensis L2TR* | 0.99 | 14.3 | 0.795 | 83 |
| AF-K0F1F1-F1-model_v4 | AFDB-PROTEOME | Hydrolase | *Nocardia brasiliensis ATCC 700358* | 0.99 | 15.5 | 0.797 | 83 |
| AF-Q9I0C5-F1-model_v4 | AFDB-PROTEOME | Chloroperoxidase | *Pseudomonas aeruginosa PAO1* | 0.99 | 16.4 | 0.792 | 85 |
| AF-K0ELX9-F1-model_v4 | AFDB-PROTEOME | Hydrolase | *Nocardia brasiliensis ATCC 700358* | 0.99 | 19.4 | 0.792 | 81 |
| AF-Q8ZLI9-F1-model_v4 | AFDB-PROTEOME | Pimeloyl-[acyl-carrier protein] methyl ester esterase | *Salmonella enterica subsp. enterica serovar Typhimurium str. LT2* | 0.99 | 18.1 | 0.792 | 82 |
| AF-A0A0H3GV49-F1-model_v4 | AFDB-PROTEOME | Putative non-heme chloroperoxidase | *Klebsiella pneumoniae subsp. pneumoniae HS11286* | 0.99 | 16.3 | 0.791 | 85 |
| AF-A0A0D2GU4 | AFDB-PROTEOM | Unplaced genomic scaffold supercont1.2, whole genome | *Fonsecaea pedrosoi CBS 271.37* | 0.99 | 15.2 | 0.789 | 85 |

| | | | | | | | |
|---|---|---|---|---|---|---|---|
| 1-F1-model_v4 | E | shotgun sequence | | | | | |
| AF-Q32AM6-F1-model_v4 | AFDB-PROTEOME | Pimeloyl-[acyl-carrier protein] methyl ester esterase | *Shigella dysenteriae Sd197* | 0.99 | 14.6 | 0.788 | 82 |
| AF-P13001-F1-model_v4 | AFDB-PROTEOME | Pimeloyl-[acyl-carrier protein] methyl ester esterase | *Escherichia coli K-12* | 0.99 | 14.6 | 0.787 | 82 |
| AF-I6YGL1-F1-model_v4 | AFDB-PROTEOME | Possible hydrolase | *Mycobacterium tuberculosis H37Rv* | 0.99 | 16.9 | 0.785 | 82 |
| AF-X8FLZ2-F1-model_v4 | AFDB-PROTEOME | Alpha/beta hydrolase fold family protein | *Mycobacterium ulcerans str. Harvey* | 0.99 | 17.5 | 0.783 | 86 |

**Table S9.** Antibacterial activities of dipyrimicin A and B reported as minimum inhibitory activity.

| | Antibacterial activity (µg/mL) | | |
|---|---|---|---|
| | Dipyrimicin A **(1)** | Dipyrimicin B **(2)** | Tetracycline (control) |
| *Bacillus spizizenii* ATCC 6633 | 32 | >128 | 0.25 |
| *Escherichia coli* MG1655 ATCC 700926 | 128 | >128 | 2 |
| *Pseudomonas aeruginosa* ATCC 27853 | >128 | NT[a] | 8 |
| *Staphylococcus aureus* ATCC 25923 | 64 | NT[a] | 0.25 |

[a]NT = not tested.

**Table S10.** ML-predicted antibacterial activity probability of BGCs encoding 2,2'-BP genes described in **Figure 6**.

| **Strain** | **NCBI RefSeq_Assembly** | ***cae* % similarity[a]** | ***col* % similarity[a]** | **Average antibacterial activity score** |
|---|---|---|---|---|
| *Actinosynnema sp.* ALI-1.44 | GCF_001984155.1_ASM198415v1_genomic | 52% | 59% | 74% |
| *Streptomyces caatingaensis* CMAA 1322 | GCF_001187435.1_ASM118743v1_genomic | 48% | 38% | 74% |
| *Amycolatopsis anabasis* EGI 650086 | GCF_009765355.1_ASM976535v1_genomic | 80% | 59% | 74% |
| *Nocardia panacis* YIM PH 21724 | GCF_003598715.1_ASM359871v1_genomic | 72% | 55% | 73% |
| *Kitasatospora albolonga* YIM 101047 | GCF_002082585.1_ASM208258v1_genomic | 68% | 88% | 70% |
| *Streptomyces sp.* SID14515 | GCF_010548505.1_ASM1054850v1_genomic | 60% | 66% | 67% |
| *Streptomyces sp.* Ncost-T6T-1 | GCF_002705685.1_ASM270568v1_genomic | 60% | 77% | 63% |
| *Streptomyces filamentosus* NRRL 15998 | GCF_000156455.1_ASM15645v1_genomic | 56% | 70% | 60% |
| *Micromonospora craniellae* LHW63014 | GCF_014764405.1_ASM1476440v1_genomic | 60% | 62% | 59% |
| *Streptomyces filamentosus* NRRL 11379 | GCF_000156695.2_ASM15669v2_genomic | 60% | 74% | 59% |
| *Streptomyces sp.* Cmuel-A718b | GCF_900092005.1_IMG- | 60% | 74% | 56% |

| | | | | |
|---|---|---|---|---|
| | taxon_2657245729_annotated_assembly_genomic | | | |
| *Micromonospora tulbaghiae* CNY-010 | GCF_003612775.1_ASM361277v1_genomic | 56% | 59% | 54% |
| *Streptomyces sp.* CS149 | GCF_003024215.1_ASM302421v1_genomic | 60% | 70% | 54% |
| *Streptomyces sp.* CAI 127 | GCF_013363655.1_ASM1336365v1_genomic | 60% | 74% | 52% |
| *Streptomyces sp.* TSRI0261 | GCF_001905485.1_ASM190548v1_genomic | 60% | 74% | 51% |
| *Streptomyces sp.* CB00271 | GCF_014495705.1_ASM1449570v1_genomic | 60% | 74% | 50% |
| *Streptomyces sp.* WA6-1-16 | GCF_020341595.1_ASM2034159v1_genomic | 60% | 74% | 49% |
| *Streptomyces sp.* CB02613 | GCF_002803175.1_ASM280317v1_genomic | 60% | 74% | 48% |

[a]from AntiSMASH 5 KnownClusterBlast
